# Evolutionary and functional divergence of Sfx, a plasmid-encoded H-NS homolog, underlies the regulation of IncX plasmid conjugation

**DOI:** 10.1101/2024.07.10.602937

**Authors:** Avril Wang, Martha Cordova, William Wiley Navarre

**Affiliations:** Department of Molecular Genetics, University of Toronto, Toronto, Ontario, Canada

## Abstract

Conjugative plasmids are widespread among prokaryotes, highlighting their evolutionary success. Despite being an important route for plasmid transmission, most natural plasmids do not constitutively conjugate and are conjugatively repressed by default. In the F plasmid, the negative regulation of conjugation is partially mediated by the chromosomal nucleoid structuring protein (H-NS). Recent bioinformatic analyses have revealed that plasmid-encoded H-NS homologs are widespread and exhibit high sequence diversity. However, the functional roles of most of these homologs and the selective forces driving their phylogenetic diversification remain unclear. In this study, we characterized the functionality and evolution of Sfx, a H-NS homolog encoded by the model IncX2 plasmid R6K. We demonstrate that Sfx, but not chromosomal H-NS, can repress R6K conjugation. Notably, we find evidence of positive selection acting on the ancestral Sfx lineage. These positively selected sites are located in the dimerization, oligomerization, and DNA-binding interfaces, many of which contribute to R6K repression activity – indicating that adaptive evolution drove the functional divergence of Sfx. We additionally show that Sfx can physically interact with various chromosomally encoded proteins, including H-NS, StpA, and Hha. Hha enhances the ability of Sfx to regulate R6K conjugation, suggesting that Sfx retained functionally important interactions with chromosomal silencing proteins. Surprisingly, the loss of Sfx does not negatively affect the stability or dissemination of R6K in laboratory conditions, reflecting the complexity of selective pressures favoring conjugation repression. Overall, our study sheds light on the functional and evolutionary divergence of a plasmid-borne H-NS-like protein, highlighting how these loosely specific DNA-binding proteins evolved to specifically regulate different plasmid functions.

**IMPORTANCE:** Conjugative plasmids play a crucial role in spreading antimicrobial resistance and virulence genes. Most natural conjugative plasmids conjugate only under specific conditions. Therefore, studying the molecular mechanisms underlying conjugation regulation is essential for understanding antimicrobial resistance and pathogen evolution. In this study, we characterized the conjugation regulation of the model IncX plasmid R6K. We discovered that Sfx, a H-NS homolog carried by the plasmid, represses conjugation. Molecular evolutionary analyses combined with gain-of-function experiments indicate that positive selection underlies the conjugation repression activity of Sfx. Additionally, we demonstrate that the loss of Sfx does not adversely affect R6K maintenance under laboratory conditions, suggesting additional selective forces favoring Sfx carriage. Overall, this work underscores the impact of protein diversification on plasmid biology, enhancing our understanding of how molecular evolution affects broader plasmid ecology.

## INTRODUCTION

Plasmids are extrachromosomal elements found across all domains of life (1–3) and can be transferred between even phylogenetically distant organisms (4). Conjugation is a major vehicle for plasmid spread, with roughly half of all fully sequenced plasmids being conjugative or mobilizable (5). Yet, most natural plasmids are conjugatively repressed (6), conjugating only with specific environmental cues or in the presence of viable recipient cells (7). Previous studies have suggested that limiting conjugation can a) offset the fitness cost of plasmid carriage by reducing energy consumption (8), b) minimize the stress response from the expression of conjugation-related genes (9, 10), and c) evade bacteriophages that attach to the conjugation pilus (11–14). Therefore, a central question in plasmid biology has been to understand the molecular mechanisms that govern conjugation repression.

Plasmid conjugation in Gram-negative bacteria depends on transfer (*tra*) genes, which encode for components involved in mating pair formation (Mpf) and DNA transfer and replication (Dtr) (15). The Mpf genes encode the type-IV secretion system (T4SS) responsible for plasmid transfer. At the same time, the Dtr proteins facilitate this process by nicking and coupling the plasmid origin of transfer (*oriT*) to the T4SS (15). Conjugation repression can occur by regulating the expression and/or activity of *tra* genes/proteins (16–22), and much of these mechanisms are gleaned from studying the model F plasmid. Repression of F plasmid conjugation involves chromosomal and plasmid-encoded factors that control *tra* gene transcription (23), translation (24), and Tra protein stability (24, 25). One such chromosomal factor is the histone-like nucleoid structuring protein (H-NS). This DNA-binding protein preferably oligomerizes along curved, AT-rich DNA, causing DNA condensation and transcriptional silencing (26, 27). Within the F plasmid, H-NS binds to *tra* gene promoters (P_Y_, P_M_, and P_J_) to repress conjugation (23, 28).

While chromosomal H-NS regulates F plasmid conjugation, many conjugative plasmids encode their own H-NS homologs. These homologs are found across different plasmid incompatibility groups, ranging from IncA/C (29), IncH (30, 31), to IncX (32, 33) – indicating a selective advantage for this alternative mode of H-NS carriage. Only a few plasmid-encoded H-NS homologs have been experimentally characterized. Sfh carried by the IncH plasmid pSfR27 is functionally redundant with chromosomal H-NS and is thought to serve as a “molecular H-NS backup (30).” Other homologs are functionally unique. Acr2 carried by the IncA/C plasmid pAR060302 specifically regulates plasmid conjugation genes without affecting most chromosomal gene expression (29). H-NS_R27_ encoded by the IncH R27 plasmid also represses conjugation and is uniquely resistant to negative regulation that typically antagonizes chromosomal H-NS (31). Across plasmid-encoded H-NS homologs, those carried by IncX plasmids (i.e., the HppX clade) are notable for their sequence diversity, especially within the typically conserved DNA-binding domain (34). While some homologs within the HppX clade are known to repress conjugation (32, 33), little is known about the processes that drove their phylogenetic diversification, the molecular mechanism of conjugation repression, their interaction with chromosomal-encoded H-NS homologs, and their broader impact on plasmid ecology.

Much of our knowledge of how H-NS regulates plasmid gene expression is gleaned from studying the F plasmid and, more recently, Sfh. However, it is unclear whether these insights are broadly applicable, especially for the phylogenetically divergent HppX clade. To address this, we characterized the evolution and function of Sfx, a H-NS homolog encoded by the model IncX2 plasmid R6K. Here, we report that the evolutionary divergence of Sfx is associated with its functional divergence from chromosomal H-NS. We find that Sfx homologs are often carried by plasmids with the MPF_T_-type T4SS and are more closely related to the H-NS paralog StpA than to H-NS. Sfx exhibits marked amino acid differences in the dimerization, oligomerization, and DNA-binding domains relative to chromosomal H-NS – many of which are driven by positive selection following the divergence of the Sfx lineage from ancestral StpA. These evolutionary signatures are accompanied by functional divergence, whereby only Sfx, but not H-NS and StpA, can repress R6K conjugation. We find that the C-terminal DNA-binding domain of Sfx is critical for its unique conjugation repression activity. In contrast, the dimerization and oligomerization domains make more subtle contributions to repression. We further show that Sfx can physically interact with chromosomal silencing factors, including H-NS, StpA, and Hha, but that only Hha contributes to Sfx-mediated conjugation repression. Additionally, we find that Sfx loss minimally impacts plasmid carrier growth rate and plasmid maintenance in laboratory culture. Collectively, our work characterizes the unique evolutionary and functional divergence of a plasmid-encoded H-NS homolog, highlighting the molecular innovation afforded by protein evolution across different genomic contexts.

## RESULTS

### Sfx is associated with mobile genetic elements and diverges from other H-NS homologs

IncX plasmids are characterized by a highly syntenic backbone, which includes a module where *sfx* is encoded immediately downstream of *topB* (DNA topoisomerase III)(35, 36). We first sought to further characterize the genomic context of Sfx homologs and their evolutionary relationships with other H-NS homologs. We queried the NCBI nr/nt database using three seed templates: Sfx, H-NS of *E. coli* K12, and StpA of *E. coli* K12. This search identified 33, 86, and 21 representative sequences using the Sfx, H-NS, and StpA seed templates, respectively (Table S1). These sequences were then aligned and used for maximum likelihood phylogenetic inference. The resultant phylogeny is divided into three main clades. Two clades consist primarily of sequences obtained using H-NS and StpA as seed templates. The third clade is composed exclusively of sequences obtained using the Sfx seed template (Figure 1A) and forms the sister group of the clade containing *E. coli* K12 StpA, consistent with previous phylogenetic analyses (34). For clarity, we designated the cluster of Sfx homologs as the Sfx clade, the group containing *E. coli* K12 H-NS as the H-NS clade, and the group containing *E. coli* K12 StpA as the StpA clade.

**Figure 1.**
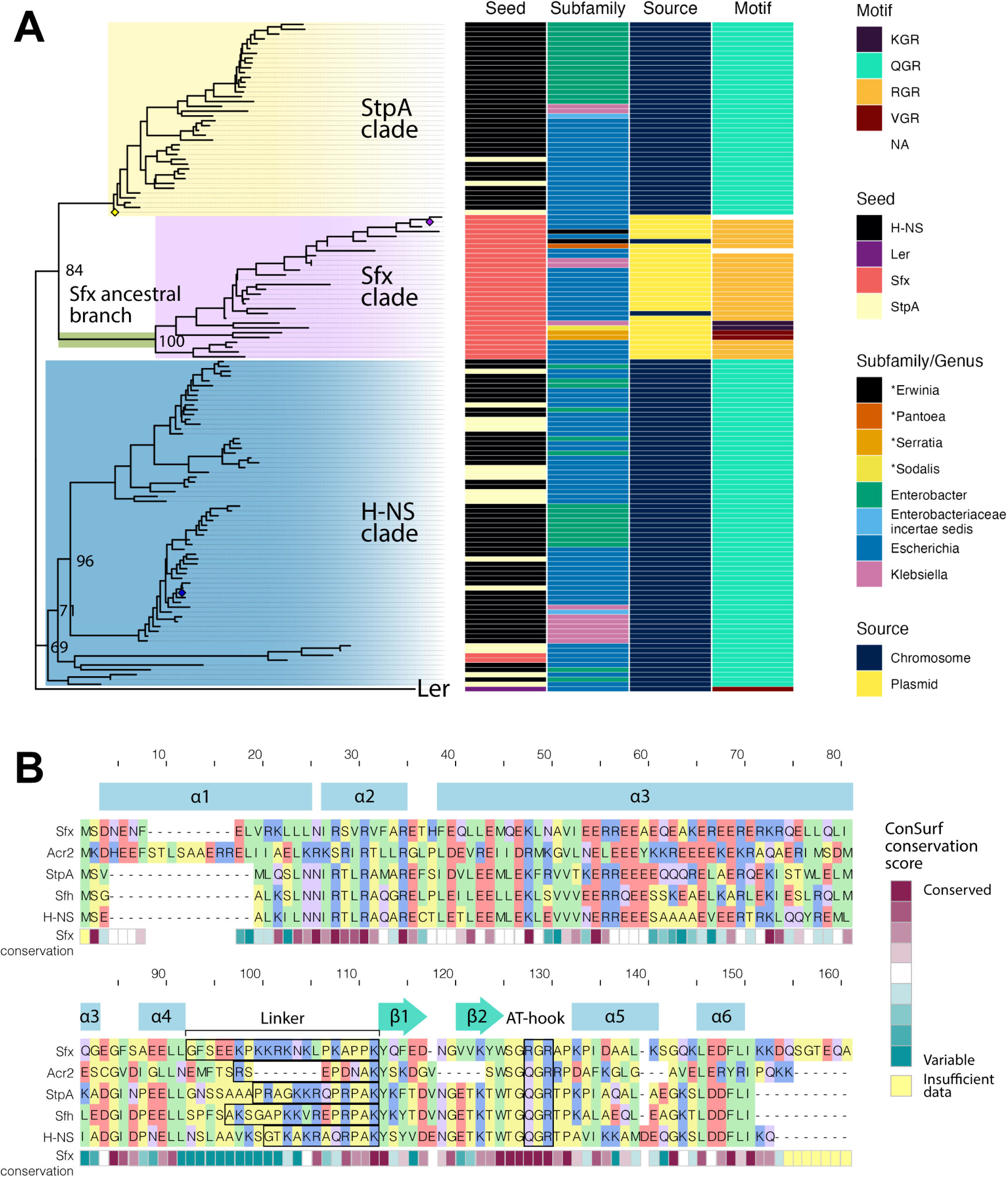
Sfx diverges phylogenetically from chromosomal and other plasmid-encoded H-NS homologs. (A) Annotated maximum likelihood phylogeny of Sfx, H-NS, and StpA homologs. Nucleotide coding sequences of the homologs are retrieved using tblastn and an in-house script. The sequences are aligned using codon-based alignment via Guidance2, and IQ-TREE2 is used to construct a maximum likelihood phylogeny. The Ler protein carried by *E. coli* O157:H7 str. Sakai, a distantly related H-NS-like protein, is used as the outgroup to root the phylogeny. Each sequence representative is annotated with associated characteristics (Seed = Template used in tblastn, Subfamily = taxonomic classification of the bacteria encoding for the protein, Source = whether the protein is located on a chromosome or plasmid, Motif = amino acid sequence of the AT-hook motif). Genera that begin with “*” do not belong to the *Enterobacteriaceae* family. The locations of Sfx, *E. coli* K12 H-NS, and *E. coli* K12 StpA are indicated by a purple, blue, or yellow diamond, respectively. The choice of foregrounds used in the Clade models is indicated and colored (StpA = yellow, Sfx = purple, H-NS = blue). The ancestral Sfx branch set as the foreground in the Branch-site model is highlighted in green. The displayed node values represent the Ultrafast Bootstrap approximates (%). (B) Sfx differs from other H-NS homologs in key residue positions. Amino acid sequences of Sfx (NCBI accession ID: WP_001282381.1), Acr2 (NCBI accession ID: WP_000651490.1), StpA (Uniprot accession: P0ACG1), Sfh (Uniprot accession: Q8GKU0), and H-NS (Uniprot accession: P0ACF8) are aligned using MAFFT G-INS-i. The secondary structure of Sfx, derived using POLYVIEW-2D of a ColabFold predicted structure, is indicated above the alignment, with α representing alpha-helices, β representing beta-sheets, and empty spaces representing coils. The degree of conservation at each residue position amongst Sfx homologs is calculated based on ConSurf analysis using default parameters. For each protein, the position of the linker, a disordered region after the fourth alpha-helix, is outlined with a black border.

Sequences from the H-NS and StpA clades are all chromosomally encoded and restricted to the *Enterobacteriaceae* family (Figure 1A). In contrast, sequences within the Sfx clade are either encoded by a plasmid or are found within chromosomal transposons (NCBI accession and range: CP013970.1:2715190-2715642 and CP028735.1:4153062-4153523). Consistent with its association with mobile genetic elements, Sfx homologs are found across a more extensive taxonomic distribution compared to proteins within the H-NS and StpA clades, including species from the *Erwinia, Pantoea, Serratia,* and *Sodalis* genera (Figure 1A).

Despite its phylogenetic clustering, the Sfx and StpA clades differ in their AT-hook motif – a region that inserts into the minor groove of AT-rich DNA and is critical for DNA binding (37). The H-NS and StpA clades and previously characterized plasmid-encoded H-NS homologs such as Acr2 and Sfh carry a QGR AT-hook motif (Figure 1A-B). In contrast, most Sfx homologs carry an RGR motif that is characteristic of xenogeneic silencing proteins encoded by bacteria with a higher genome GC content, including Lsr2 from *Mycobacteria* (38) and Bv3F from *Burkholderia vietnamiensis* (39). Another distinct property of Sfx is its longer and more proline-rich predicted linker (4 prolines in Sfx linker vs. 1-3 prolines in other H-NS-like proteins; Figure 1B), which could increase coil rigidity, limit the formation of larger nucleoprotein complexes, and increase target selectivity (40). Collectively, our findings indicate that Sfx homologs form a phylogenetically distinct clade associated with mobile genetic elements. This evolutionary divergence is partly driven by differences in residues that could impact protein oligomerization and DNA binding.

### Sfx homologs are often carried by plasmids with the MPF_T_ T4SS

Given the association between Sfx homologs and plasmids, we next explored its distribution across 55 representative IncX plasmid lineages (Table S2). Despite significant sequence divergence among the plasmid lineages — with no identifiable core ORFs according to Roary (41) — a little more than half of the plasmids (31/55; 56.4%) encode for H-NS homologs. Most of these plasmids carry a H-NS homolog with the RGR AT-hook motif (25/31; 80.6%) characteristic of the Sfx clade (Figure S1). Furthermore, plasmid-encoded H-NS and the MPF_T_/VirB-like T4SS tend to co-occur (Figure S1). Conversely, plasmids lacking a MPF system or possessing a MPF_F_ T4SS (i.e., the type of T4SS possessed by the F plasmid) tend not to carry Sfx or any other H-NS homolog (Figure S1). Taken together, our findings reaffirm the conservation of Sfx across IncX plasmids and reveal a close association of Sfx with MPF_T_ T4SS, suggesting that Sfx may act as a regulator uniquely specific for MPF_T_-type conjugation systems.

### Positive selection drove the evolutionary divergence of Sfx

Next, we used molecular evolution analysis to investigate the evolutionary processes that drove the divergence of the Sfx clade. We first employed Clade Models to determine the selective forces affecting extant sequences. Our best-performing model (CmD with Sfx and StpA clades set as separate foregrounds) suggests that most protein sites of extant H-NS, StpA, and Sfx clades show signatures of purifying selection (Table S3, Figure S2), indicating their functional importance. Notably, the StpA clade exhibits the least degree of selective constraints (Figure S2), which may reflect its recessive phenotype (42) and indicate its potential for adaptive evolution.

We next used the Branch-Site test to determine the evolutionary processes that drove the divergence of the ancestral Sfx lineage from the StpA clade (ancestral branch highlighted in green in Figure 1A). The Branch-Site test detects signatures of positive selection from a single branch selected *a priori* (the foreground) within a phylogeny (the background). Adaptive evolution/positive selection can be inferred if fitting an alternative Branch-Site model incorporating positive selection leads to a statistically significant improvement in model fit relative to fitting a null model, where only neutral and purifying evolution is considered (43). We find that fitting the alternative Branch-Site model leads to a statistically significant (*p-*value = 0.015) improvement in model fit relative to the null model (Table S3), suggesting that positive selection occurred after the Sfx clade diverged from ancestral StpA.

We next analyzed the potential functionality of positively selected residues indicated by the Naive Empirical Bayes (NEB) and Bayes Empirical Bayes (BEB) analyses (see Table 1 for details). While NEB outputs are more prone to false positives than BEB for small datasets (44), both methods produced similar results (Figure 2A; Table 1). Most positively selected sites are conserved within the Sfx, H-NS, and StpA clades (Figure S3), indicating their functional importance. Overall, positive selection altered the dimerization and oligomerization interfaces of Sfx relative to H-NS (Figure 2B), which may change the protein’s nucleation profile and heteromeric interactions. For instance, the N9L mutation (corresponding to the L15 (Sfx)-N9 (H-NS) transition) reduces H-NS’ capacity to interact with Hha and repress the hemolysin (*hly*) operon (45), indicating that Sfx may display altered interaction with Hha. Although not directly part of the dimerization interface, the T27-C21 transition is biochemically similar to the C21S mutation in H-NS, also known to decrease DNA-binding affinity (46). Changes to the oligomerization domain may also impact protein regulation. H-NS is temperature sensitive, with higher temperature destabilizing the oligomerization interface (e.g., by disrupting the ionic bond D71:R54’) (47, 48), causing H-NS to adopt an inhibited configuration (47) and relieve repression (49–51). In Sfx, positive selection led to divergence in many of the sites involved in H-NS dimer-dimer interaction and H-NS auto-inhibition (e.g., S77-D71; Figure 2B, Table 1) (47, 48).

**Figure 2.**
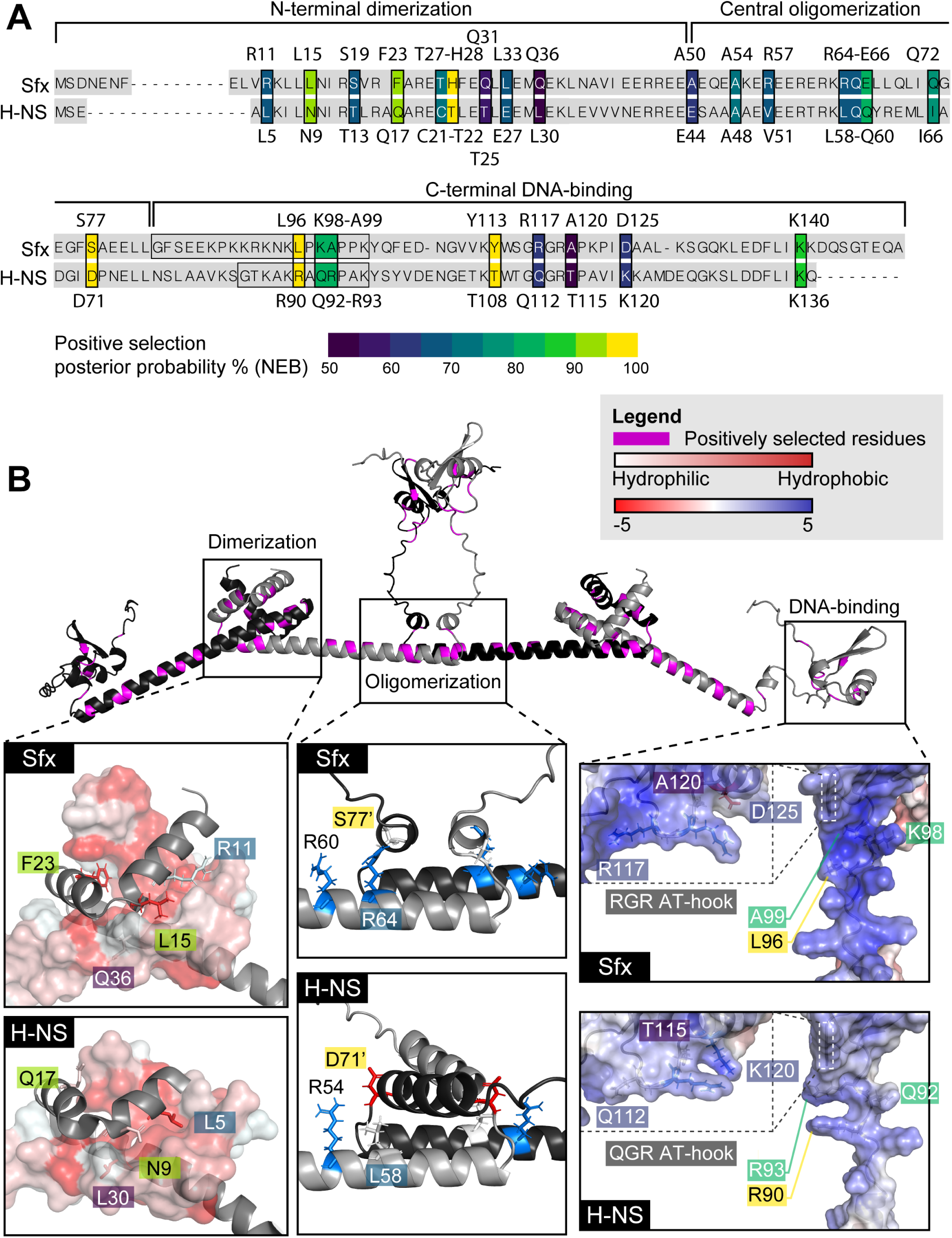
Positively selected sites map to Sfx dimerization, oligomerization, and DNA-binding interfaces. (A) Positions of positively selected sites mapped onto an alignment of Sfx (R6K) and *E. coli* K12 H-NS. The identity and position of each positively selected site are displayed above (Sfx)/below (H-NS) the alignment. Each site is colored based on the posterior probability that the site has experienced positive selection following the divergence of the ancestral Sfx lineage (Naive Empirical Bayes results). The linker region of each protein is outlined with a black box. (B) Positively selected sites mapped onto the predicted structures of Sfx and H-NS. The top panel displays the ColabFold-predicted tetrameric structure of Sfx. Residues that have undergone positive selection are colored in magenta. The interaction interfaces of the dimerization, oligomerization, and DNA-binding domains are displayed in the inset boxes, with the top panel containing the predicted Sfx structure and the bottom panel containing the predicted H-NS structure. The N-terminal dimerization domain inset displays the interaction between a dimer (one as a grey cartoon structure and the other as a surface representation structure). The hydrophobicity of the surface representation structure is indicated by its color, and positively selected residues that form the hydrophobic interaction interface of H-NS are labeled. The central oligomerization domain inset displays the interaction between two dimers (one colored in grey and the other colored in black). The locations and charge (blue = positively charged, red = negatively charged) of positively selected residues participating in H-NS oligomerization are labeled. The apostrophe indicates that the residue is located on another protein. Though not positively selected, R60/R54 (Sfx/H-NS) is highlighted due to its ionic interaction with D71 in H-NS. The DNA-binding domain inset displays a surface representation of the linker region and the AT-hook motif (highlighted by a dashed box) colored based on its electrostatic charge (blue = positively charged, red = negatively charged). The locations of positively selected residues that participate in DNA binding in H-NS are labeled.

**Table 1.**
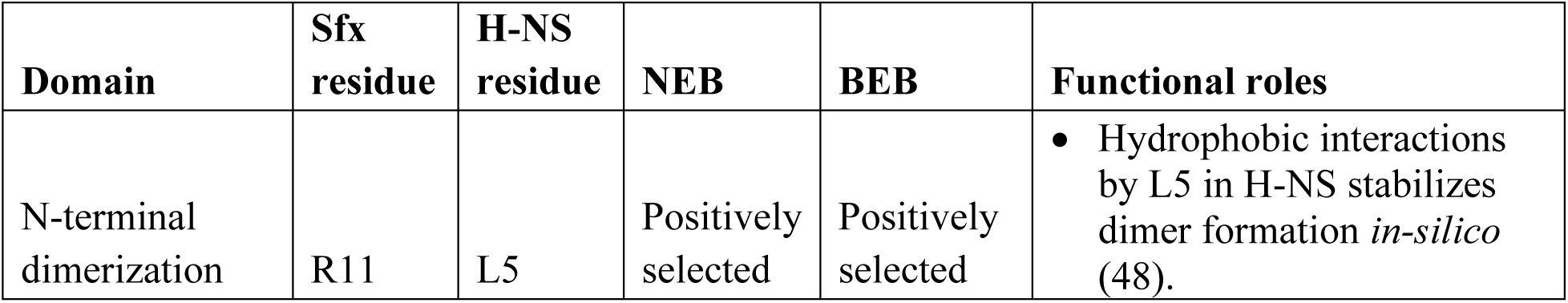
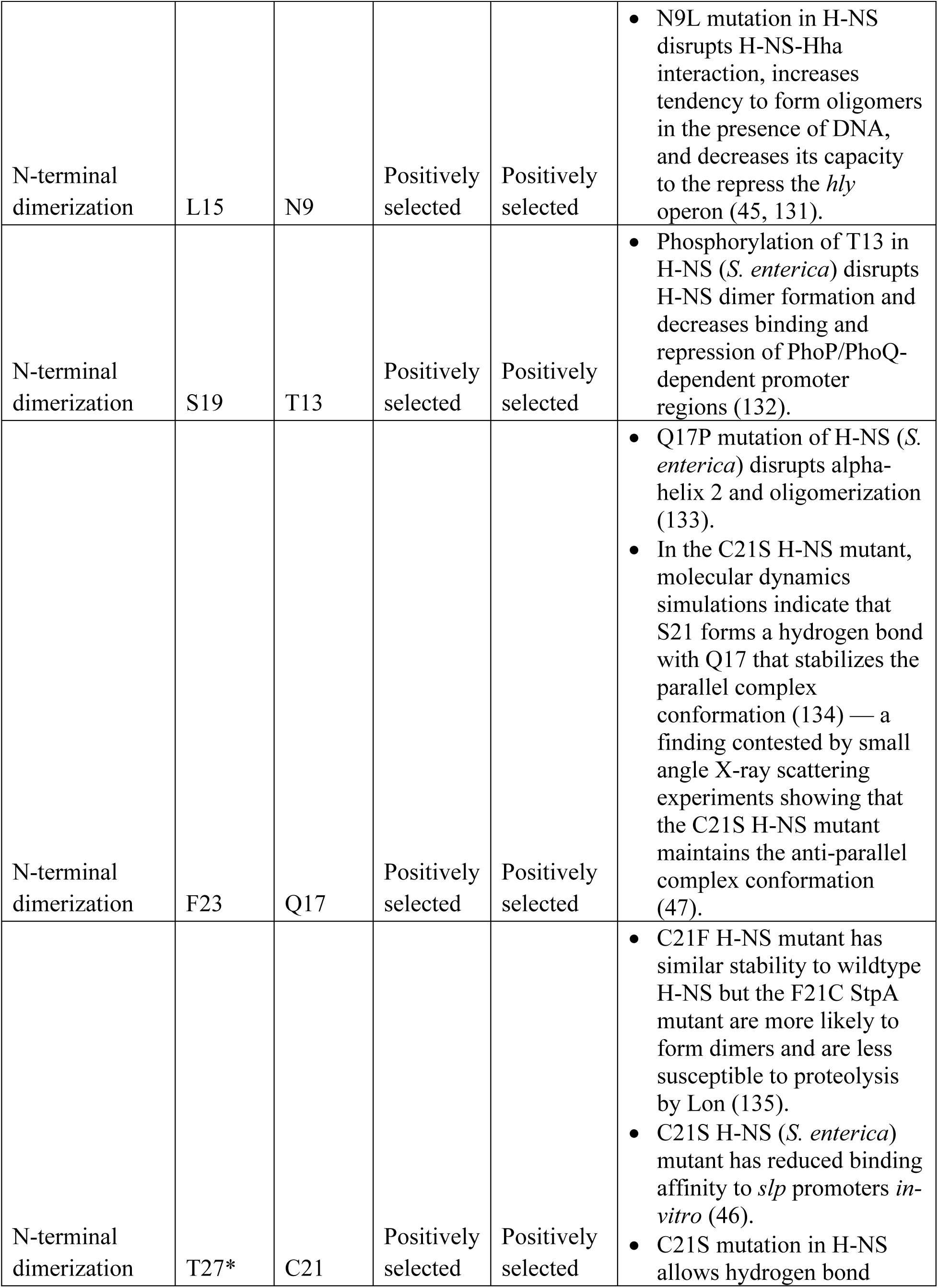
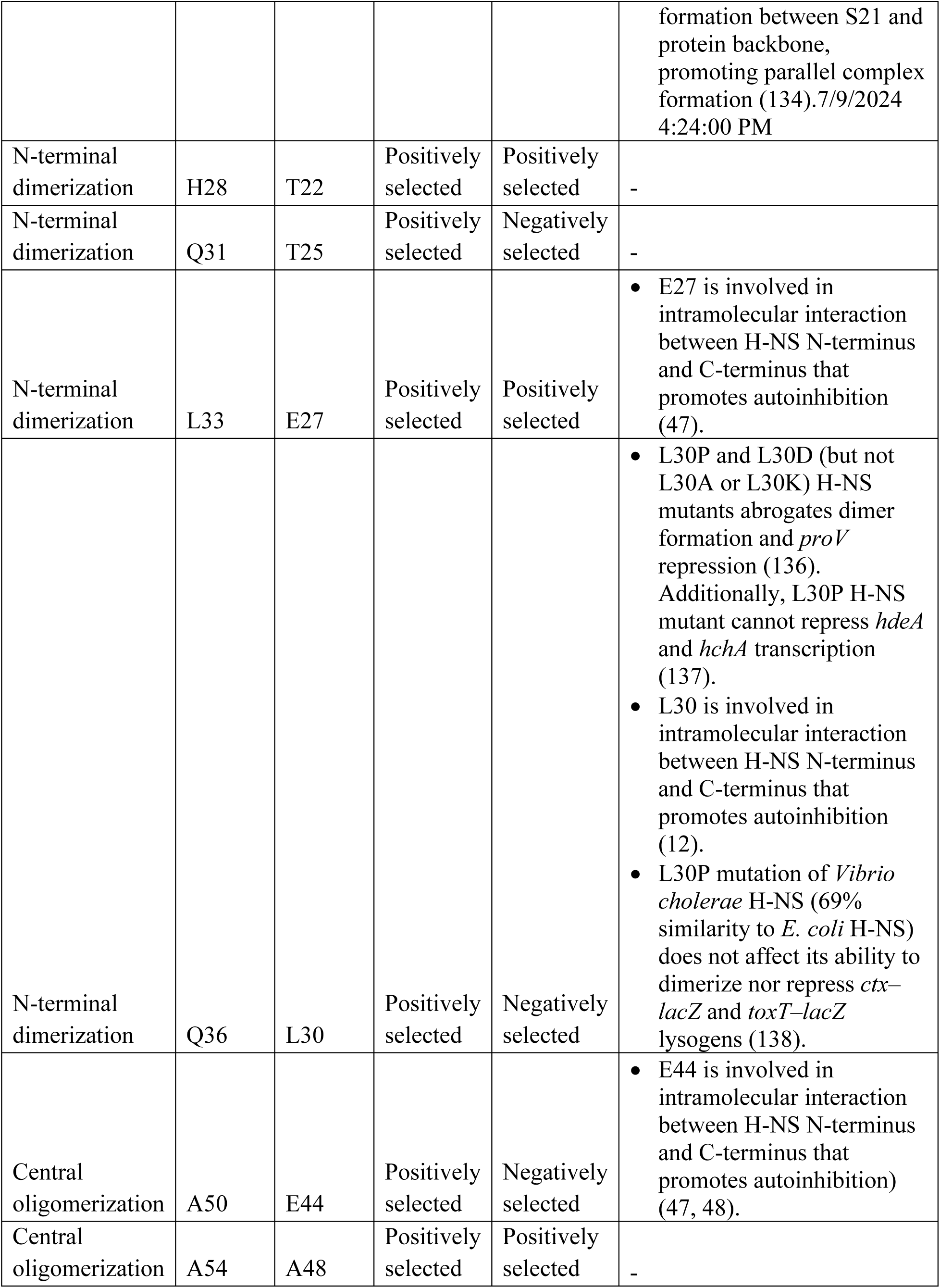
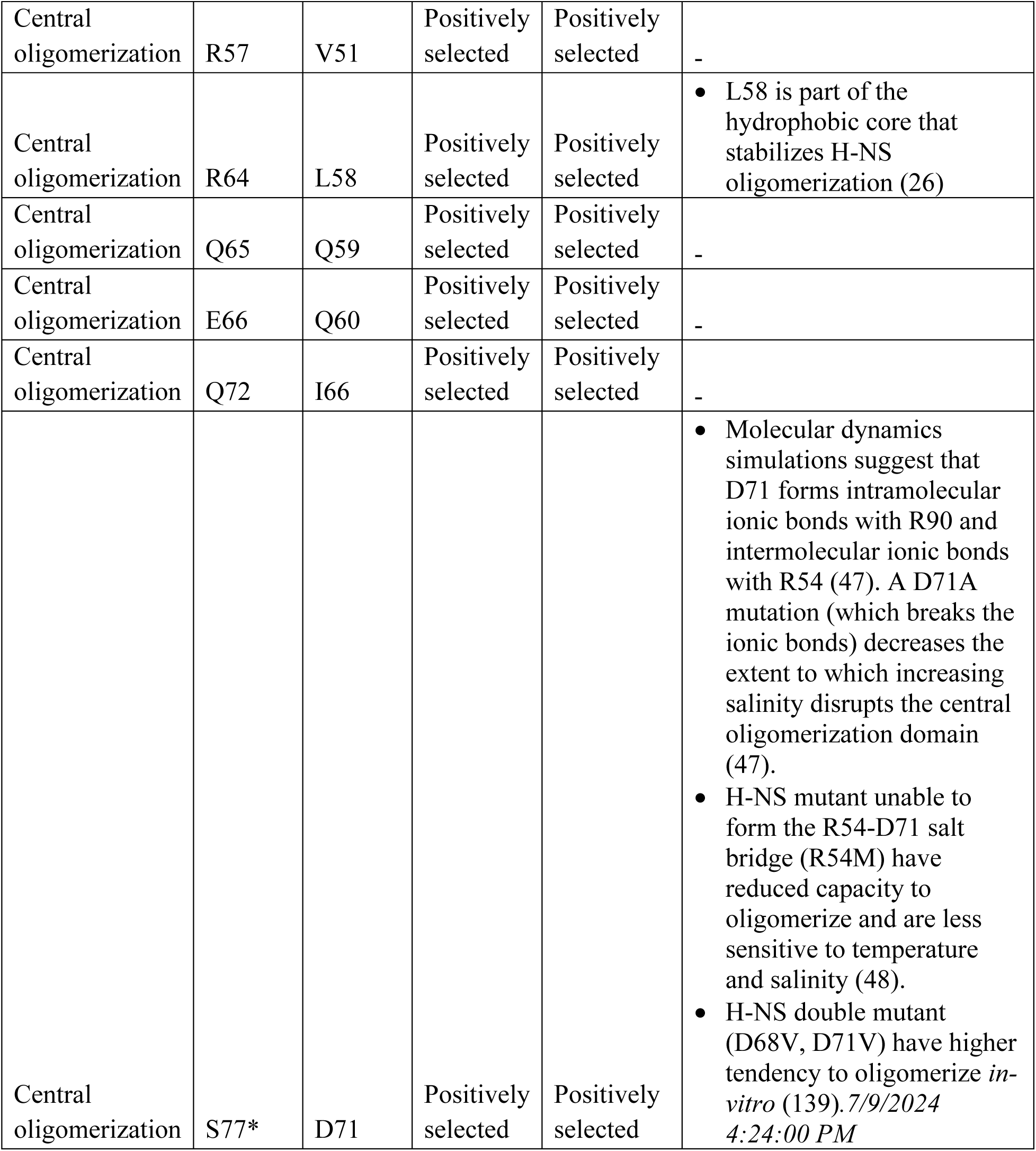
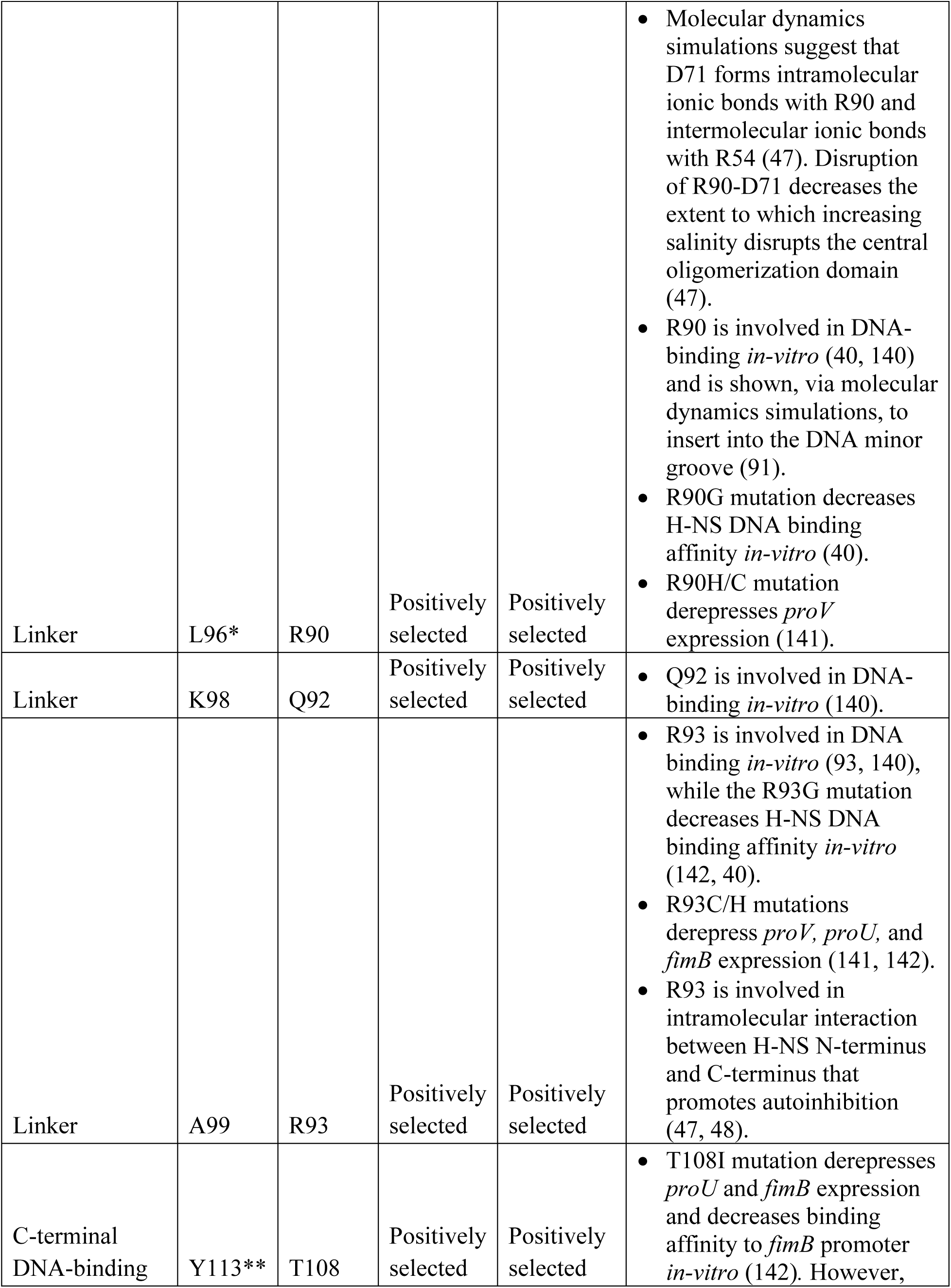
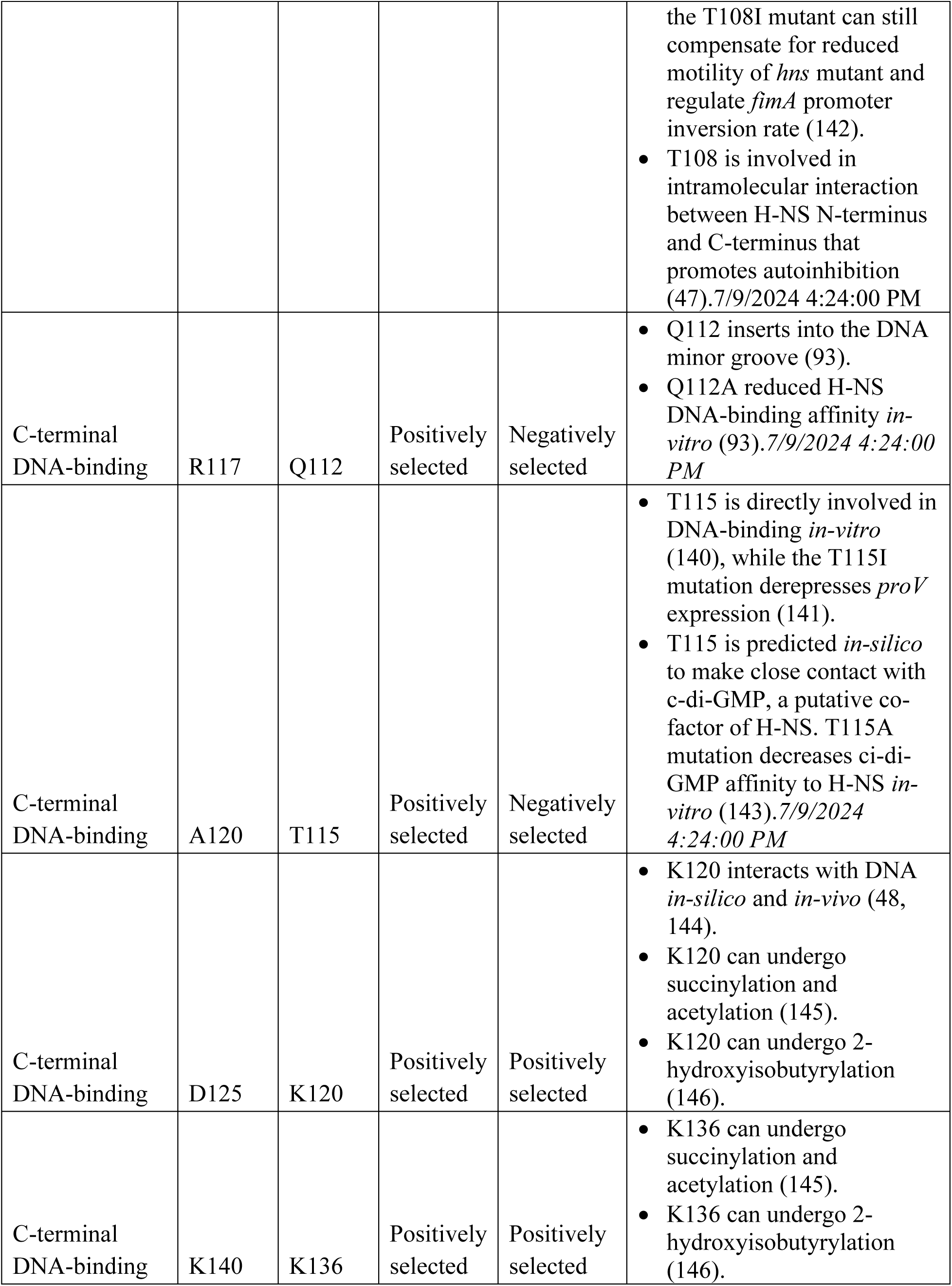
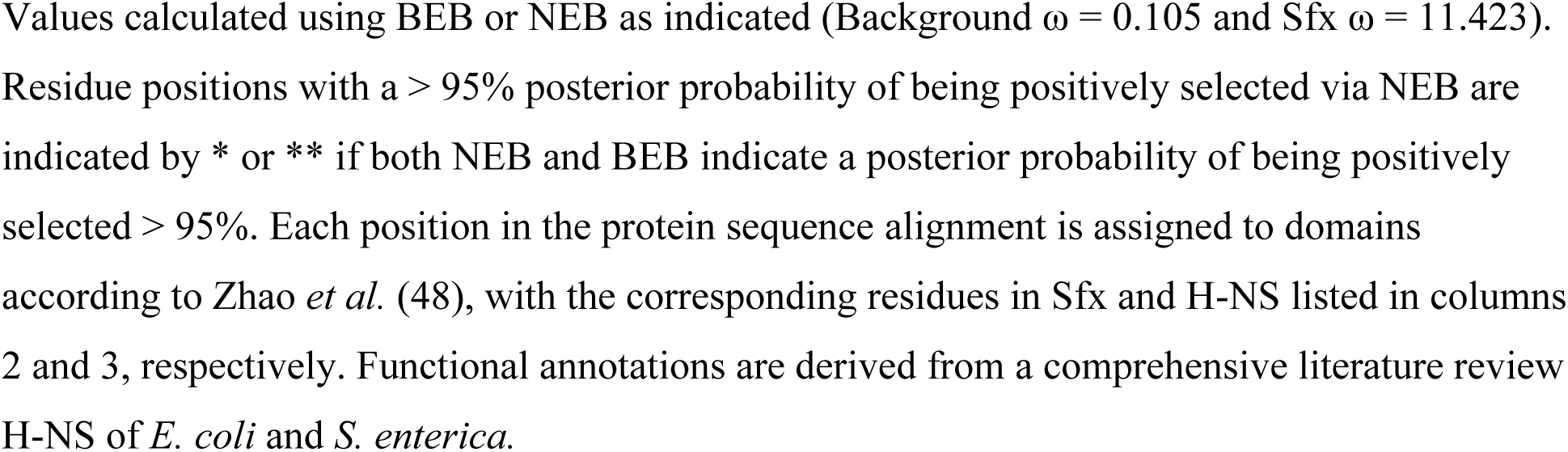
Characteristics of positively selected residues defined based on NEB and BEB analyses.

| Domain | Sfx residue | H-NS residue | NEB | BEB | Functional roles |
| --- | --- | --- | --- | --- | --- |
| N-terminal dimerization | R11 | L5 | Positively selected | Positively selected | Hydrophobic interactions by L5 in H-NS stabilizes dimer formation *in-silico* (48). |
| N-terminal dimerization | L15 | N9 | Positively selected | Positively selected | • N9L mutation in H-NS disrupts H-NS-Hha interaction, increases tendency to form oligomers in the presence of DNA, and decreases its capacity to repress the *hly* operon (45, 131). |
| N-terminal dimerization | S19 | T13 | Positively selected | Positively selected | • Phosphorylation of T13 in H-NS (*S. enterica*) disrupts H-NS dimer formation and decreases binding and repression of PhoP/PhoQ-dependent promoter regions (132). |
| N-terminal dimerization | F23 | Q17 | Positively selected | Positively selected | • Q17P mutation of H-NS (*S. enterica*) disrupts alpha-helix 2 and oligomerization (133).<br>• In the C21S H-NS mutant, molecular dynamics simulations indicate that S21 forms a hydrogen bond with Q17 that stabilizes the parallel complex conformation (134) — a finding contested by small angle X-ray scattering experiments showing that the C21S H-NS mutant maintains the anti-parallel complex conformation (47). |
| N-terminal dimerization | T27* | C21 | Positively selected | Positively selected | • C21F H-NS mutant has similar stability to wildtype H-NS but the F21C StpA mutant are more likely to form dimers and are less susceptible to proteolysis by Lon (135).<br>• C21S H-NS (*S. enterica*) mutant has reduced binding affinity to *slp* promoters *in-vitro* (46).<br>• C21S mutation in H-NS allows hydrogen bond<br>formation between S21 and protein backbone, promoting parallel complex formation (134).7/9/2024 4:24:00 PM |
| N-terminal dimerization | H28 | T22 | Positively selected | Positively selected | - |
| N-terminal dimerization | Q31 | T25 | Positively selected | Negatively selected | - |
| N-terminal dimerization | L33 | E27 | Positively selected | Positively selected | E27 is involved in intramolecular interaction between H-NS N-terminus and C-terminus that promotes autoinhibition (47). |
| N-terminal dimerization | Q36 | L30 | Positively selected | Negatively selected | L30P and L30D (but not L30A or L30K) H-NS mutants abrogates dimer formation and *proV* repression (136). Additionally, L30P H-NS mutant cannot repress *hdeA* and *hchA* transcription (137).<br>L30 is involved in intramolecular interaction between H-NS N-terminus and C-terminus that promotes autoinhibition (12).<br>L30P mutation of *Vibrio cholerae* H-NS (69% similarity to *E. coli* H-NS) does not affect its ability to dimerize nor repress *ctx-lacZ* and *toxT-lacZ* lysogens (138). |
| Central oligomerization | A50 | E44 | Positively selected | Negatively selected | E44 is involved in intramolecular interaction between H-NS N-terminus and C-terminus that promotes autoinhibition (47, 48). |
| Central oligomerization | A54 | A48 | Positively selected | Positively selected | - |
| Central oligomerization | R57 | V51 | Positively selected | Positively selected | - |
| Central oligomerization | R64 | L58 | Positively selected | Positively selected | • L58 is part of the hydrophobic core that stabilizes H-NS oligomerization (26) |
| Central oligomerization | Q65 | Q59 | Positively selected | Positively selected | - |
| Central oligomerization | E66 | Q60 | Positively selected | Positively selected | - |
| Central oligomerization | Q72 | I66 | Positively selected | Positively selected | - |
| Central oligomerization | S77* | D71 | Positively selected | Positively selected | • Molecular dynamics simulations suggest that D71 forms intramolecular ionic bonds with R90 and intermolecular ionic bonds with R54 (47). A D71A mutation (which breaks the ionic bonds) decreases the extent to which increasing salinity disrupts the central oligomerization domain (47).<br>• H-NS mutant unable to form the R54-D71 salt bridge (R54M) have reduced capacity to oligomerize and are less sensitive to temperature and salinity (48).<br>• H-NS double mutant (D68V, D71V) have higher tendency to oligomerize *in-vitro* (139).<br>7/9/2024 4:24:00 PM |
| Linker | L96* | R90 | Positively selected | Positively selected | • Molecular dynamics simulations suggest that D71 forms intramolecular ionic bonds with R90 and intermolecular ionic bonds with R54 (47). Disruption of R90-D71 decreases the extent to which increasing salinity disrupts the central oligomerization domain (47).<br>• R90 is involved in DNA-binding *in-vitro* (40, 140) and is shown, via molecular dynamics simulations, to insert into the DNA minor groove (91).<br>• R90G mutation decreases H-NS DNA binding affinity *in-vitro* (40).<br>• R90H/C mutation derepresses *proV* expression (141). |
| Linker | K98 | Q92 | Positively selected | Positively selected | • Q92 is involved in DNA-binding *in-vitro* (140). |
| Linker | A99 | R93 | Positively selected | Positively selected | • R93 is involved in DNA binding *in-vitro* (93, 140), while the R93G mutation decreases H-NS DNA binding affinity *in-vitro* (142, 40).<br>• R93C/H mutations derepress *proV*, *proU*, and *fimB* expression (141, 142).<br>• R93 is involved in intramolecular interaction between H-NS N-terminus and C-terminus that promotes autoinhibition (47, 48). |
| C-terminal DNA-binding | Y113** | T108 | Positively selected | Positively selected | • T108I mutation derepresses *proU* and *fimB* expression and decreases binding affinity to *fimB* promoter *in-vitro* (142). However,<br>the T108I mutant can still compensate for reduced motility of *hns* mutant and regulate *fimA* promoter inversion rate (142).<br>• T108 is involved in intramolecular interaction between H-NS N-terminus and C-terminus that promotes autoinhibition (47). |
| C-terminal DNA-binding | R117 | Q112 | Positively selected | Negatively selected | • Q112 inserts into the DNA minor groove (93).<br>• Q112A reduced H-NS DNA-binding affinity *in-vitro* (93). |
| C-terminal DNA-binding | A120 | T115 | Positively selected | Negatively selected | • T115 is directly involved in DNA-binding *in-vitro* (140), while the T115I mutation derepresses *proV* expression (141).<br>• T115 is predicted *in-silico* to make close contact with c-di-GMP, a putative co-factor of H-NS. T115A mutation decreases ci-di-GMP affinity to H-NS *in-vitro* (143). |
| C-terminal DNA-binding | D125 | K120 | Positively selected | Positively selected | • K120 interacts with DNA *in-silico* and *in-vivo* (48, 144).<br>• K120 can undergo succinylation and acetylation (145).<br>• K120 can undergo 2-hydroxyisobutyrylation (146). |
| C-terminal DNA-binding | K140 | K136 | Positively selected | Positively selected | • K136 can undergo succinylation and acetylation (145).<br>• K136 can undergo 2-hydroxyisobutyrylation (146). |

Values calculated using BEB or NEB as indicated (Background $\omega = 0.105$ and Sfx $\omega = 11.423$ ). Residue positions with a $> 95\%$ posterior probability of being positively selected via NEB are indicated by \* or \*\* if both NEB and BEB indicate a posterior probability of being positively selected $> 95\%$ . Each position in the protein sequence alignment is assigned to domains according to Zhao *et al.* (48), with the corresponding residues in Sfx and H-NS listed in columns 2 and 3, respectively. Functional annotations are derived from a comprehensive literature review H-NS of *E. coli* and *S. enterica*.

Among H-NS homologs, only the HppX clade significantly deviates from chromosomal H-NS at the DNA-binding domain (34). Our data suggests that positive selection drove the transitions of key residues that directly interact with DNA in H-NS (Figure 2B; Table 1), many of which are positively charged. Interestingly, we find that positive selection underlies the unique RGR AT-hook motif of Sfx (Figure 2B; only under positive selection by NEB but not BEB), further supporting its potential functional significance. Together, our results suggest that the ancestral Sfx lineage experienced positive selection at numerous sites, many of which correspond to residues implicated in protein-protein interaction, environmental regulation, and DNA binding.

### Sfx, but not H-NS and StpA, represses R6K conjugation

We next examined whether the sequence divergence displayed by Sfx is of functional importance. To test this, we performed filter-paper conjugation experiments involving *E. coli* carrying wildtype R6K plasmid or one lacking *sfx* (R6KΔ*sfx*). Deletion of *sfx* increased R6K conjugation efficiency on filter-paper by >1000-fold (Figure 3A). Considering that the filter-paper is an artificial setting for conjugation, we examined whether Sfx loss would also increase R6K spread in a more naturalized biofilm setting (52). We find that R6K*Δsfx* conjugates at a higher efficiency than R6K in biofilms (∼5-fold; Figure 3B), affirming that Sfx is a negative regulator of plasmid transmission.

**Figure 3.**
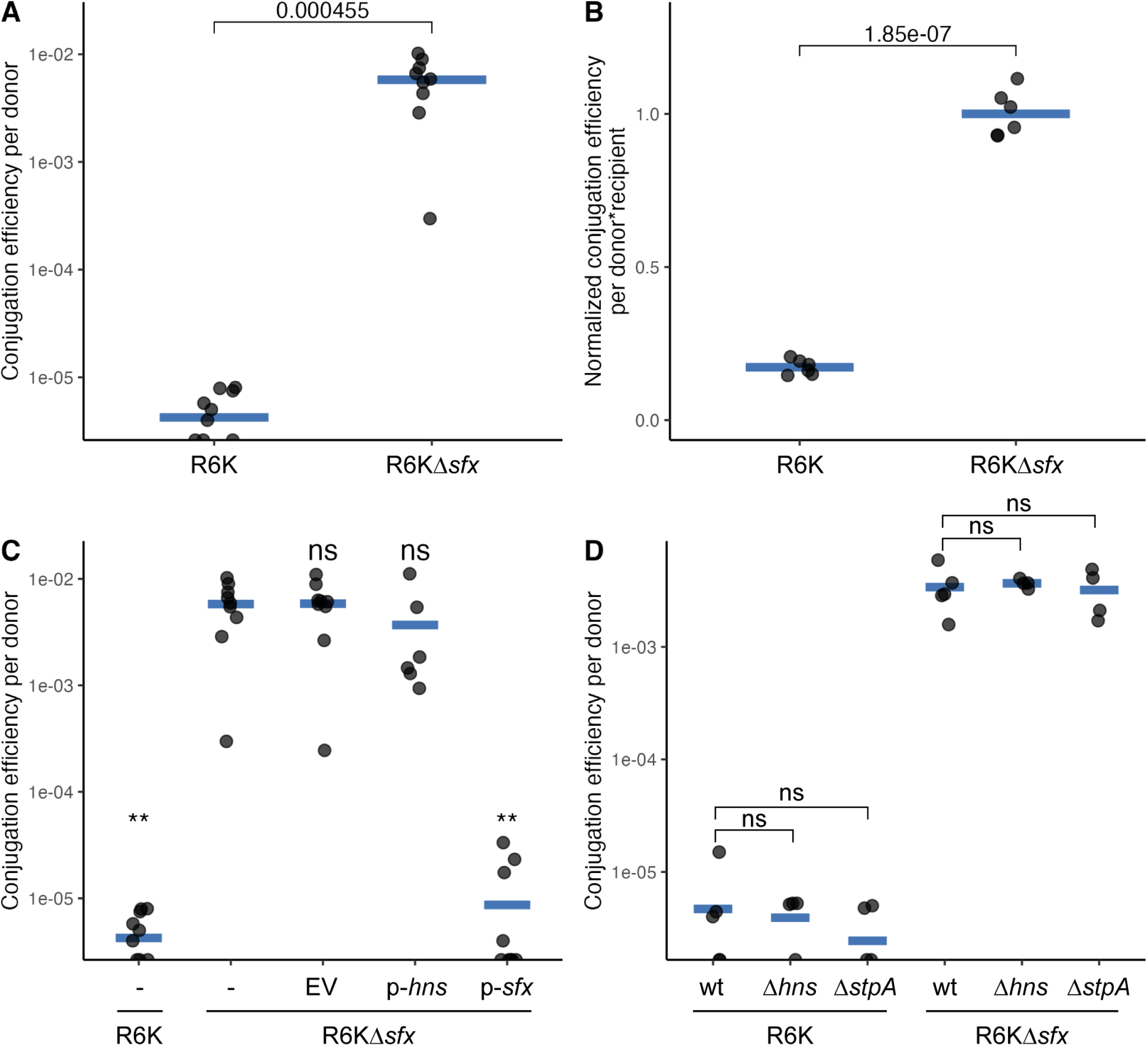
Sfx represses R6K conjugation on filter-paper and within biofilms. (A) *sfx* deletion increases R6K conjugation on filter-paper. Two OD_600_U of donor (*E. coli* K12 carrying R6K or R6KΔ*sfx*) and recipient cells (NaN_3_^r^ EcoR25) are incubated on a filter-paper disc at 30°C for 3 hours. Conjugation efficiency is measured by the number of transconjugants divided by the number of donors at the end of the conjugation period. (B) *sfx* deletion increases R6K conjugation in biofilm. Donor (*E. coli* K12 carrying R6K or R6KΔ*sfx*) and recipient cells (NaN_3_^r^ EcoR25) are seeded in M9+0.2% glucose media to a final OD_600_ of 0.03 and grown at 30°C for 24 hours. Conjugation efficiency is assessed by dividing the number of transconjugant cells by the product of donor and recipient populations (to account for potential differences in recipient population growth). Conjugation efficiency is further normalized by scaling the mean conjugation efficiency of R6KΔ*sfx* carriers to 1. (C) Sfx, but not H-NS, complementation decreases R6KΔ*sfx* conjugation efficiency. The y-axis displays the conjugation efficiency of different donor cells (NaN_3_^r^ EcoR25 recipient cells) on filter-paper. The x-axis indicates the plasmids carried by the donor cells. (D) Chromosomal H-NS and StpA do not repress R6K conjugation. The y-axis displays the conjugation efficiency of different donor cells on filter-paper. The x-axis indicates the donor’s genotype (top row) and plasmid (bottom row). (A-D) The data displayed are pooled results from at least two experimental replicates, and each point represents one biological replicate (averaged value across two technical replicates). The Student’s t-test is used to assess statistical significance, and the Benjamini-Hochberg correction is applied when more than one t-test is conducted. Abbreviations: ns (*p-*value ≥ 0.05); - (placeholder); EV (pHSG576 empty vector, low-copy); p*-hns* (pHSG576-*hns,* native *hns* promoter); p-*sfx* (pHSG576-*sfx,* native *sfx* promoter).

To ensure that the impact of Sfx loss is not polar, we transformed a separate low-copy plasmid expressing Sfx from its native promoter into donor cells. We find that complementation reduced the conjugation efficiency of R6KΔ*sfx* donors to levels comparable to wildtype R6K donors (Figure 3C). Interestingly, expressing chromosomal H-NS from the same low-copy plasmid backbone does not affect the conjugation efficiency of R6KΔ*sfx* (Figure 3C). To test whether chromosomal H-NS and its paralog StpA can also regulate R6K conjugation, we repeated the filter-paper conjugation experiments using Δ*hns* and Δ*stpA* donors. We find that neither of these chromosomal deletions affect the conjugation efficiency of R6K and R6KΔ*sfx* (Figure 3D), indicating that chromosomal H-NS homologs do not regulate R6K conjugation. Collectively, our data suggest that Sfx represses R6K conjugation and has functionally diverged from chromosomal H-NS and StpA.

### Sfx can partially compensate for H-NS loss

Plasmid-encoded H-NS homologs H-NS_R27_ and Sfh can partially compensate for Δ*hns* phenotypes (53, 54). Considering H-NS’s inability to repress R6K conjugation, we wondered whether Sfx, in contrast, can rescue the null phenotypes of *E. coli Δhns*. The presence of R6K or a low-copy plasmid expressing Sfx from its native promoter compensates for the reduced motility and growth lag of *hns* mutants (55) (Figure 4A, C). Conversely, while R6KΔ*sfx* carriage does not significantly impact the growth dynamics of wildtype *E. coli* (Figure 4A), it reduces the growth rate of *hns* mutants (Figure 4D), suggesting that at least one H-NS homolog (chromosomal H-NS or Sfx) is needed to mitigate the fitness burden of R6K carriage. Accordingly, expression of H-NS or Sfx from its native promoter off a low-copy plasmid rescued the growth defect exhibited by the *hns* mutant carrying R6K*Δsfx* (Figure 4E). Overall, while H-NS cannot repress R6K conjugation, Sfx can partially compensate for H-NS function, suggesting a partial, albeit asymmetrical, overlap between the regulons of H-NS and Sfx.

**Figure 4.**
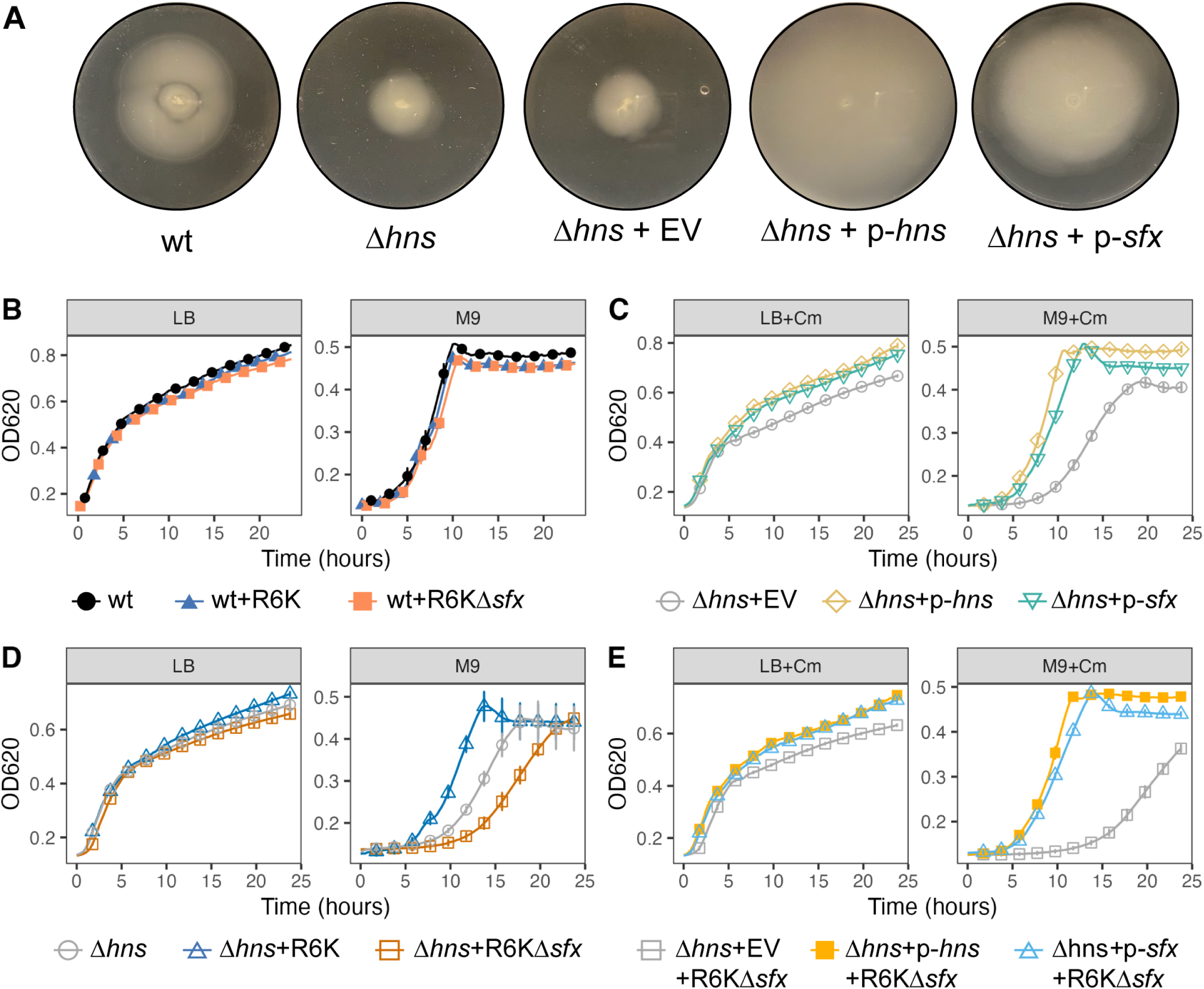
Sfx can partially compensate for Δ*hns* phenotypes. (A) Expressing Sfx increases the motility of *hns* mutants. Motility assays are performed by inoculating 6 µL of *E. coli* Keio Collection strains (genotype displayed below the plates) subcultured to a mid-log phase (OD_600_∼0.6) into the center of a soft-agar plate (0.3% agar). The plates are incubated at 30°C for 24 hours. The images are representative of two independent experimental replicates, each with two biological replicates. (B-E) display time-series growth curve data of various *E. coli* strains grown at 37°C in LB or M9+0.2% glucose. When required, 20 µg/mL of chloramphenicol (“Cm”) is added to maintain stable inheritance of the pHSG576 low-copy plasmids. Each data point represents the mean ± standard deviation across two biological replicates, each with two technical replicates. The data are representative of three independent experimental replicates. Abbreviations: wt (wildtype *E. coli* BW25113); EV (pHSG576 empty vector, low-copy); p*-hns* (pHSG576-*hns,* native *hns* promoter); p-*sfx* (pHSG576-*sfx,* native *sfx* promoter).

### The DNA-binding domain is necessary for Sfx-mediated conjugation repression

Given that Sfx functionally diverges from H-NS, we performed a series of gain-of-function experiments to explore which structural elements confer Sfx’s unique ability to repress R6K conjugation. We constructed chimeric proteins with different combinations of H-NS and Sfx domains (indicated in Figure 5A), expressed them from a low-copy plasmid off the native *sfx* promoter, and assessed their impact on the conjugation efficiency of R6KΔ*sfx*.

**Figure 5.**
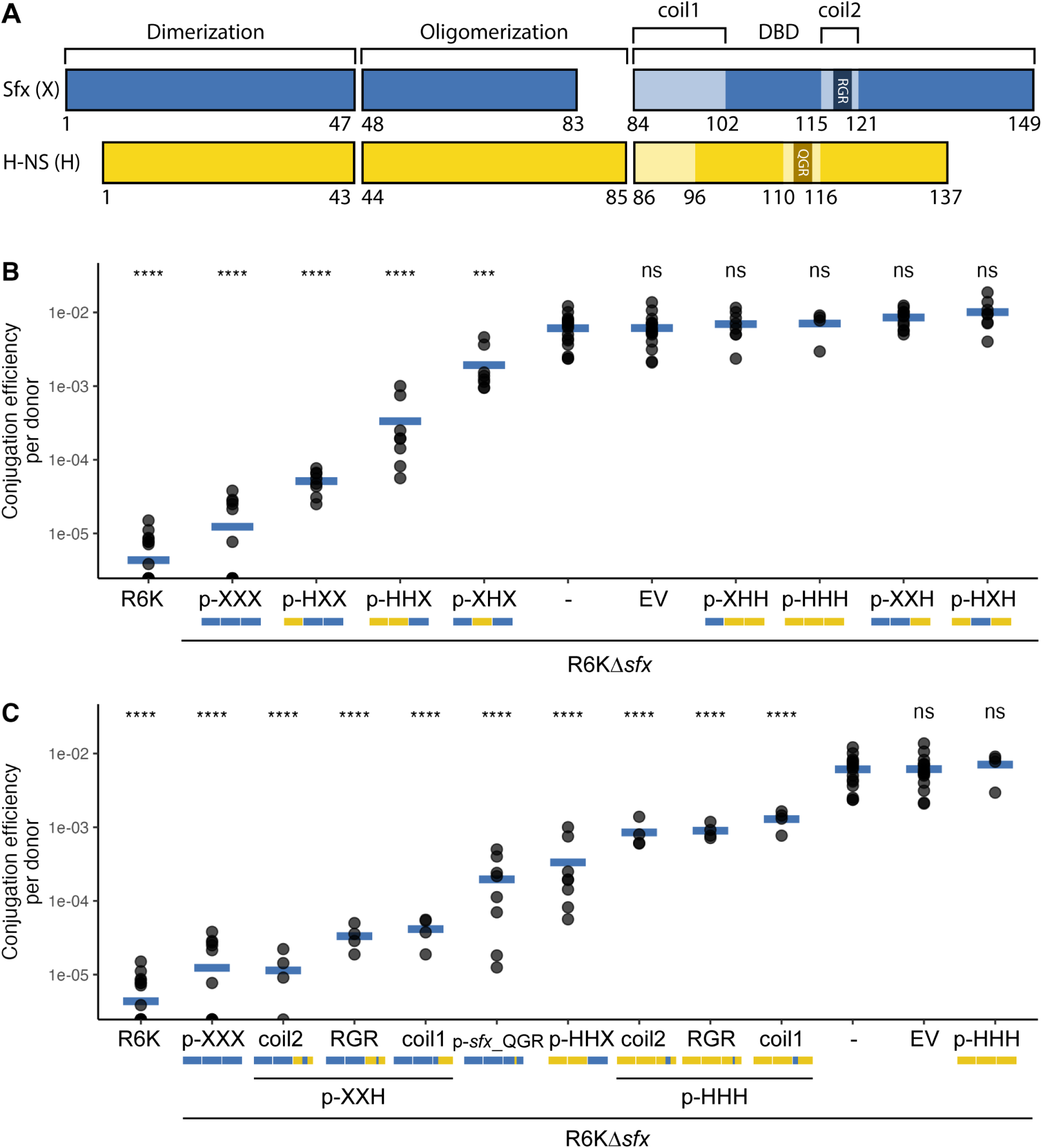
Sfx C-terminus is necessary for R6K conjugation repression. (A) Schematic of domain assignment for Sfx and H-NS. Secondary structure assignments are based on the ColabFold-predicted structure of Sfx and H-NS. Coil 1 (linker) and coil 2 (contains the AT-hook motif) are two unstructured regions within the DNA-binding domain (DBD). The precise amino acid position at which the domains are split (and where the chimeras are fused) is displayed below each protein schematic. (B-C) display the results from filter-paper conjugation experiments performed with various donors (indicated on the x-axis) and NaN_3_^r^ EcoR25 recipient. The identity of the chimeric proteins expressed from a low-copy plasmid (*sfx* promoter) is represented by p-NNN where each N represents the domain identities (X = Sfx, H = H-NS) and a mini graphic schematic (blue = Sfx, yellow = H-NS). (B) The Sfx DNA-binding domain is needed for R6K conjugation repression. The y-axis displays the conjugation efficiency (transconjugant population divided by donor population) of donors carrying R6K or R6KΔ*sfx* and low-copy plasmids encoding for chimeric proteins. Only R6KΔ*sfx* carriers that express chimeric proteins with the Sfx DNA-binding domain (p-NNX) exhibit statistically significantly reduced (*p-*value < 0.05) conjugation efficiency relative to no plasmid control (R6KΔ*sfx* only). (C) The linker and the AT-hook motif contribute to Sfx’s conjugation repression activity. The same conjugation assay is performed as in (B), albeit with donors expressing chimeric proteins with point or segmental mutations. Coil 1/2 indicates chimeric proteins carrying Sfx’s coil 1/2 (in either a p-XXH or p-HHH background), whereas p-*sfx_*QGR represents plasmid encoding for a Sfx protein with the QGR AT-hook motif. For (B-C), statistical significance is assessed using the Student’s t-test (reference group is *E. coli* carrying only R6KΔ*sfx*) with Benjamini-Hochberg correction. The data are representative results from two independent experimental replicates. Abbreviations: **** (adjusted *p-*value < 0.0001); *** (adjusted *p-*value < 0.001); ns (adjusted *p-*value ≥ 0.05).

All chimeras containing Sfx’s DNA-binding domain can repress conjugation of R6KΔ*sfx* to varying degrees (Figure 5B), suggesting that the DNA-binding domain is important for Sfx’s repression activity. The addition of Sfx’s central oligomerization domain improves chimeric protein’s repression activity (p-HXX vs. p-HHX; Figure 5B), which could suggest that the oligomerization domain contributes to Sfx repression only when the C-terminal domain is present and/or that the presence of the domain improves protein expression/folding.

To narrow down the regions within Sfx’s DNA-binding domain that contribute to conjugation repression, we swapped smaller segments of the H-NS DNA-binding domain with corresponding segments from Sfx. Exchanging either the linker region (coil 1) or the coiled region surrounding the AT-hook motif (coil 2) conferred partial conjugation repression to constructs containing the H-NS DNA binding domain (Figs 5B and 5C), suggesting that Sfx’s linker and its unique RGR AT-hook motif are major contributors to its unique repression activity. To further confirm that the functional importance of coil 2 is derived from the RGR AT-hook motif (and not the surrounding residues), we constructed Sfx mutants that carried the QGR AT hook-motif and H-NS mutants that carried the RGR AT hook-motif. The RGR to QGR mutation decreased but did not completely abolish Sfx’s conjugation repression activity (p-*sfx_*QGR; Figure 5C), suggesting that the RGR AT-hook motif and other motifs (e.g., linker) function independently to repress R6K conjugation. Strikingly, the H-NS mutant bearing a single QGR to RGR mutation can partially repress R6KΔ*sfx* conjugation with an efficiency similar to that of H-NS chimeras carrying the entire Sfx coil 2 (∼10-fold repression, Figure 5C). These findings indicate that the RGR AT-hook motif alone accounts for the functional importance of coil 2. Altogether, our results provide evidence that the sequence variations displayed within the DNA-binding domain of Sfx are central to its unique conjugation repression activity.

### Sfx interacts with Hha to repress R6K conjugation

H-NS regulatory activity is modulated by its interactions with StpA (42, 56, 57), Hha (58, 49, 57), and Cnu/YdgT (59, 60). StpA is a paralog of H-NS that can interact with H-NS to form more thermally stable heterodimers (61) and bridged filaments (H-NS:StpA filaments that connect two DNA duplexes) (62, 57). These H-NS:StpA filaments promote greater RNA polymerase pausing (57) and could be important for gene regulation at high temperatures where H-NS repression is relieved (49, 50). H-NS also interacts with Hha and its paralog Cnu via its dimerization domain (63, 64). H-NS-Hha interaction promotes the formation of bridged filaments that stimulate greater RNA polymerase pausing (57) and is needed to repress expression of various virulence-related genes (e.g., *hlyCABD* operon (49, 58), LEE pathogenicity island (65, 66), *Salmonella* pathogenicity island 2 (67)) The role of Cnu is less understood (60), though recent *in-vitro* work suggests that Cnu may be involved osmolarity response (68).

Given the importance of protein-protein interactions for H-NS activity and the fact that plasmid-encoded H-NS, such as Sfh, are known to interact with chromosomal H-NS and StpA (69), we used bacterial-two-hybrid to examine the physical interaction between Sfx, H-NS, StpA, Hha, and Cnu. Due to the toxicity of H-NS overexpression in the presence of ampicillin (70), we used a truncated H-NS lacking its DNA-binding domain (M1-G85) for assays involving high-copy vectors (Amp^R^) encoding for H-NS.

We first validated our assay by recovering previously characterized interactions, including those between H-NS and StpA (71), H-NS and Hha (72), H-NS and Cnu (59, 73), and StpA and Cnu (73) (Figure 6A). We find that Sfx interacts with H-NS, StpA, and Hha, while Sfx-Cnu interaction is only detected when Cnu is expressed from a high-copy vector (Figure 6A-B). Interestingly, fusion of the reporter fragment to the N- or C-terminus of Hha did not affect Hha-Sfx interaction, unlike Hha-H-NS interaction that is weakened by Hha C-terminal fusion (Figure 6A) (72). This discrepancy in Hha sensitivity to terminal fusions indicates potential differences between Sfx-Hha and H-NS-Hha interactions, which corroborates with the fact that key residues involved in H-NS-Hha interactions underwent positive selection in the ancestral Sfx lineage (e.g., L15-N9 transition).

**Figure 6.**
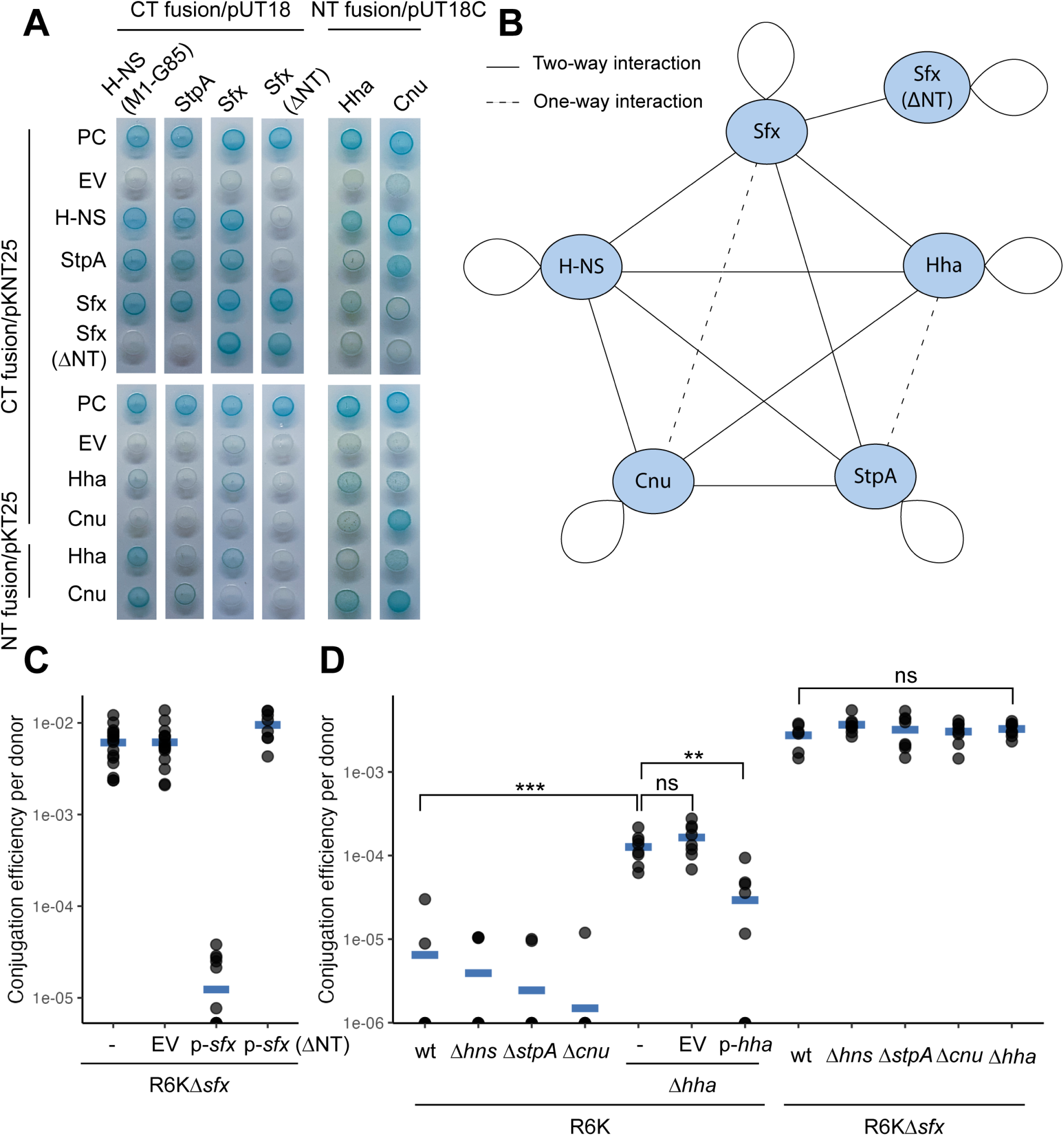
Sfx-Hha interaction contributes to conjugation repression. (A) Sfx interacts with H-NS, StpA, and Hha. Bacterial-two-hybrid assay is used to assess binary protein-protein interactions between H-NS, Sfx, StpA, Hha, and Cnu. A blue spot indicates that the expressed fusion proteins interact with sufficient strength to reconstitute the split adenylate cyclase, leading to expression of a cAMP-CAP-regulated *lacI.* A white spot indicates either insufficient interaction between the two fusion proteins or insufficient expression. Fusion proteins are either made by fusing the adenylate cyclase fragment to the C-terminus (CT) or N-terminus (NT). H-NS (M1-G85) lacking the DNA-binding domain is used in lieu of the full-length protein due to the toxicity of H-NS expression from a high-copy vector in the presence of ampicillin. Sfx (ΔNT) represents an N-terminally truncated variant (ΔD3-E8). Each assay is performed with a positive control (PC; *E. coli* carrying pKNT25-*hns* and pUT18-*stpA –* two proteins that are known to interact) and a negative control (*E. coli* carrying the plasmid indicated on the top row and a pKNT25/pKT25 empty vector). The assay is performed with three biological replicates, each with two technical replicates. The presented images are representative of two independent experiments. (B) Network depiction of bacterial-two-hybrid results. The solid lines suggest that protein interaction is detected for both protein construct combinations (e.g., pUT18-*hns* and pKNT25-*stpA*, pUT18-*stpA* and pKNT25-*hns*). The dotted lines suggest that protein interaction is detected only with one of the protein construct combinations. (C) N-terminally truncated Sfx cannot repress R6K conjugation. Various constructs (“-” = no plasmid, EV = empty vector, p-*sfx* = pHSG576-*sfx*, p-*sfx* (ΔNT) = pHSG576-*sfx* (ΔE3-D8)) are transformed into *E. coli* carrying R6KΔ*sfx* and tested for their ability to repress conjugation efficiency on filter-paper. (D) Hha co-represses R6K conjugation with Sfx. Filter-paper conjugation is performed with various *E. coli* donors from the Keio collection. The genotypes of the donors are indicated by the top row in the x-axis, and the plasmids they carry are displayed in the bottom row. Complementation of *hha* deletion is performed by transforming R6K carriers with EV (pHSG576 empty vector) or p-*hha* (pHSG576-*hha* with native *hha* promoter, low-copy). For (C-D), the data shown is representative of 2 independent experiments. When indicated, the Student’s t-test with Benjamini-Hochberg correction is used to assess statistical significance. Abbreviations: *** (adjusted *p-*value < 0.001); ** (adjusted *p-*value < 0.01); ns (adjusted *p-*value ≥ 0.05).

Interestingly, Sfx that lacks its N-terminal extension (ΔD3-E8/ΔNT) cannot form heteromeric interactions but can still interact with wildtype Sfx and N-terminally truncated Sfx (Figure 6A). N-terminally truncated Sfx cannot repress R6KΔ*sfx* conjugation when expressed from a low-copy plasmid (Figure 6C), suggesting that heteromeric interactions are needed for Sfx-mediated conjugation repression and/or that the N-terminal extension is necessary for proper Sfx-Sfx interaction. To explore the former possibility, we assessed whether Sfx’s interactions with H-NS, StpA, Hha, and Cnu are necessary for its conjugation repression activity. To this end, we performed filter-paper conjugation using R6K/R6KΔ*sfx-*carrying donors with *hns, stpA, hha,* or *cnu* deletions (gene deletions verified by lack of PCR amplification). We find that *hns, stpA,* and *cnu* deletions do not affect the conjugation efficiency of R6K and R6KΔ*sfx* (Figure 6D). In contrast, *hha* deletion increases the rate of R6K conjugation by ∼10-fold, and this elevated conjugation efficiency is decreased by the complementation of Hha encoded on a low-copy vector (Figure 6D). Moreover*, hha* deletion does not elevate the conjugation rate of R6KΔ*sfx*, indicating that Hha repression of R6K conjugation is dependent on Sfx (Figure 6D). This result and our bacterial-two-hybrid findings indicate that Sfx may interact with Hha to repress R6K conjugation.

### Sfx loss does not affect R6K fitness in laboratory settings

Conjugation is an energetically expensive process that can sometimes decrease the fitness of plasmid carriers (74, 75). We have so far been unable to detect a fitness cost (i.e., growth defect in wildtype *E. coli*) associated with carrying R6KΔ*sfx* in liquid culture (Figure 4B), which is inconsistent with the expected tradeoff between conjugation and growth (76) and the widespread prevalence of Sfx homologs across MPF_T_-type IncX plasmids (Figure S1). Therefore, we next investigated whether this lack of fitness cost of Sfx loss also applies in biofilms, a conjugative-permissive environment where plasmid fitness is dictated by donor (vertical inheritance) and transconjugant (horizontal inheritance) population dynamics. When co-cultured with EcoR25 recipient cells, we find that R6K and R6KΔ*sfx* donors reached a similar population density in the biofilm and planktonic states (Figure S4A). This result is affirmed by an R6K/R6KΔ*sfx* competition assay, where both plasmid carriers reached a similar population density in the biofilm and planktonic states (Figure S4B). Overall, we cannot detect a fitness cost to Sfx loss in a laboratory biofilm setting.

We next investigated whether Sfx loss would incur a fitness cost on a longer timescale. To test this, we serially passaged 6 independent strains of *E. coli* BW25113 carrying R6K or R6KΔ*sfx* in LB or minimal M9+0.2% glucose media for 20 days (∼200 generations). Under both media conditions, all lineages stably maintained R6K and R6KΔ*sfx* (Figure 7A), suggesting that Sfx loss does not impact plasmid stability within the study period – consistent with previous studies of an IncX3 plasmid (33). To investigate if compensatory mutations on the plasmids could account for this stability, we performed whole plasmid sequencing of the ancestor and two independent evolved lineages for each plasmid and passaging condition (a total of four evolved R6K and four R6KΔ*sfx* sequenced). No mutations were detected on the plasmids after 20 days of serial passaging. To investigate whether the passaging selected for chromosomal mutations that affected plasmid conjugation efficiency, we selected two colonies (R6KΔ*sfx* carriers) from each lineage and passaging condition and compared their conjugation rate relative to their ancestors. We find that passaging in LB and M9 media did not impact the conjugation efficiency of R6KΔ*sfx* carriers (one lineage passaged in LB displayed slightly higher conjugation efficiency; Figure 7B), suggesting that short-term passaging did not select for lower conjugation efficiency. This data is consistent with our inability to detect a fitness cost associated with Sfx loss.

**Figure 7.**
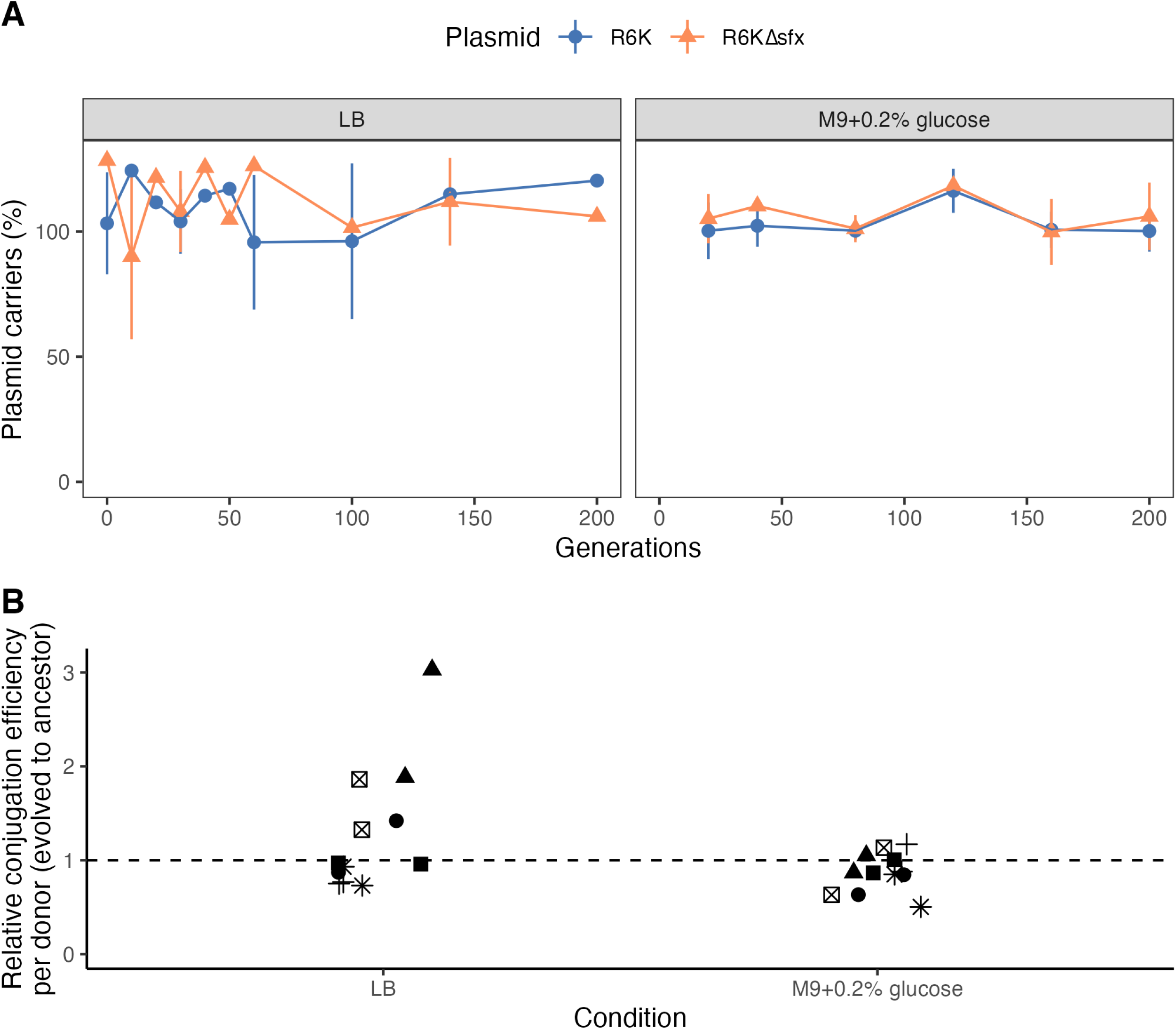
Sfx does not affect R6K stability. (A) R6K and R6KΔ*sfx* are stably maintained throughout serial passaging in LB and M9 media. For each passaging experiment, 6 independent lineages are diluted 1:000 in fresh media without antibiotics every 24 hours. The proportion of plasmid carriers is assessed by dividing the number of streptomycin-resistant colonies by the total colony count. Each point represents the mean proportion of plasmid carriers, and the lines delineate the mean ± standard deviation. (B) Passaging in LB or M9 does not decrease donor conjugation efficiency. The filter-paper conjugation experiment is performed using the ancestral (day 0) and evolved (day 20) strains from the passaging experiment shown in (A). The relative conjugation efficiency of the evolved-to-ancestor strain is plotted on the y-axis. The different shapes indicate the lineage of each sample, and each point represents the value of a biological replicate (averaged across two technical replicates).

## DISCUSSION

Plasmid conjugation is ubiquitous in natural bacterial communities and is crucial in driving bacterial evolution (77–79, 1). Interestingly, conjugation is repressed by default across most natural conjugative plasmids (6), indicating widespread selective pressures that drove the convergent evolution of repression systems. Understanding the molecular mechanism and evolution of conjugation repression is of major interest in plasmid biology. Here, we report that the evolutionary divergence of Sfx, a plasmid-encoded H-NS homolog, contributes to the unique regulation of IncX plasmid conjugation.

Gene duplication has long been recognized as a major driver for genome evolution (80, 81). Duplicated genes, known as paralogs, provide functional redundancy that temporarily relaxes purifying selection, allowing for sequence divergence that leads to gene pseudogenization (becoming non-functional), conservation (maintaining the same function), subfunctionalization (performing a subset of roles of the ancestral gene), or neofunctionalization (performing a new function) (81). Our results suggest that this duplication-divergence framework may explain the evolution of Sfx and the unique regulatory system of some IncX plasmids.

First, our phylogenetic analysis indicates that Sfx homologs form the sister clade of the H-NS paralog StpA (Figure 1A), consistent with prior work (34). This relationship suggests that ancestral Sfx may also be a H-NS paralog and likely experienced reduced purifying selection following the duplication. Indeed, our molecular evolution analysis suggests that the extant StpA clade experiences more relaxed selective constraints relative to other H-NS homologs (Figure S2), presumably due to the lack of a pronounced fitness cost associated with *stpA* null mutations (42, 82). We propose that this recessive phenotype of StpA, and likely other H-NS paralogs, potentiates the evolution of functionally distinct H-NS proteins.

Both neutral evolution (83) and positive selection (84–86) are known to drive sequence divergence following gene duplication. What drove the divergence of ancestral Sfx following its duplication? Our evolutionary analyses suggest that positive selection contributed to the initial burst of evolution experienced by the ancestral Sfx lineage (Figure 2). However, neutral evolution dominated in the clade’s later evolution (Table S3), implying the loss of functional redundancy following lineage divergence. We propose that this later neutral evolution of the Sfx clade may be facilitated by the proliferation of homologs across IncX plasmids, particularly those with the MPF_T_-type T4SS – with plasmid incompatibility curbing gene recombination and a higher gene copy number potentially boosting mutation rates (87, 88).

Having established the mechanism of Sfx clade evolution, we next explored the functional consequences of these mutations. Indeed, many of the positively selected sites we document align with H-NS regions important for dimerization, oligomerization, interaction with Hha, DNA binding, and environmental sensing (Table 1). Some of these variations underpin the functional divergence between Sfx and chromosomal H-NS. Notably, Sfx, but not H-NS nor StpA, can repress conjugation of R6K (Figure 3D) – a feature that likely favored the migration/persistence of ancestral Sfx onto/on IncX plasmids. The C-terminal DNA-binding domain of Sfx is necessary for conjugation repression. Additionally, Sfx-mediated repression is enhanced by its interaction with chromosomal Hha and requires Sfx’s N-terminal dimerization domain (Figure 6A). This co-repression may involve Hha’s ability to promote DNA bridging and generate topological stress (89, 57), inhibiting *tra* gene transcription and/or prohibiting the initial unwinding of the donor strand needed for conjugative transfer. Conversely, plasmid-encoded topoisomerase III (*topB*), which frequently colocalizes with *sfx* (35, 36), could relieve this topological stress and promote conjugation. Given that similar Hha-H-NS co-repression is observed in the IncHI plasmid R27 (53) and that Hha homologs are often co-localized with H-NS homologs in plasmids (90), conjugation regulation by H-NS/Hha pairs may be a common mechanism.

After demonstrating the functional importance of Sfx’s C-terminal DNA-binding domain, we further explored the biochemical basis of its significance. Our gain-of-function experiments indicate that the linker region is important for Sfx’s conjugation repression activity (Figure 5C). Positive selection within the linker region notably redistributed its positive charge (Figure 2B). While the functional impact of this charge variation is unclear — since similar changes in chromosomal H-NS did not significantly impact its regulatory activity (91) — it is plausible that the distinct charge distribution, longer length, and increased proline-induced rigidity (40) of the Sfx linker could collectively contribute to its unique repression activity.

The RGR AT-hook motif is another distinct feature of the Sfx clade (Figure 1A) and contributes to the regulatory activity of Sfx’s C-terminal DNA binding domain (Figure 5C). The first position of this motif displays signatures of positive selection (Figure 2A). Remarkably, altering just one amino acid from QGR to RGR in chromosomal H-NS enables it to partially repress R6K conjugation (Figure 5C), demonstrating that small sequence changes can confer large functional shifts. Interestingly, the RGR AT-hook motif is typically found in distantly related H-NS-like proteins encoded by bacteria with GC-rich genomes. These include Lsr2 carried by *Mycobacteria* (62-70% GC) (92) and Bv3F carried by *Burkholderia* (39) (59.2%-68.9% GC; from NCBI genome). These H-NS-like proteins show a heightened affinity for higher GC% sequences than those with the QGR motif (39, 93), enabling them to better differentiate between core and foreign genes in high GC% genomic contexts (93, 94).

Curiously, Sfx homologs are not found in high GC% genomic contexts (Figure S5B). For instance, R6K has a GC% of 45.3%, slightly lower than *E. coli* BW25113 (50.8%; NCBI accession: CP009273.1). Furthermore, IncX plasmids carrying Sfx homologs display an unusual base composition: their insertion elements (foreign cargo genes) are higher in GC% than their core genes (median GC% difference of 7.4%; Figure S5A-B), unlike previously characterized plasmids (95, 96). We hypothesize that Sfx’s RGR AT-hook motif and the plasmid’s atypical GC composition may facilitate better regulation of plasmid conjugation. First, the *tra* operon of R6K is less AT-rich than typical H-NS binding sites (56.6% AT from *taxA* to *tiv11* vs. 61.4% AT for average H-NS-bound site (97)). Therefore, the RGR motif may enable Sfx to bind and repress transcription more effectively in these regions, where the higher GC content prohibits efficient regulation by chromosomal H-NS. This bias towards Sfx-mediated regulation, rather than H-NS-mediated, could confer several selective advantages, including optimized timing of conjugation independent of H-NS activity and adaptability to hosts without H-NS homologs. The elevated GC% of insertion elements also suggests potential decoupling of insertion element regulation from conjugation, enabling appropriate cargo gene expression that enhances plasmid carrier fitness. Overall, Sfx represents a unique instance of an RGR-bearing H-NS homolog found in a lower GC% genomic context. Future experiments examining the exact mechanism of how the RGR AT-hook motif facilitates conjugation regulation could yield invaluable insights.

Conjugation repression is common among characterized plasmids (6), and the fitness cost of constitutive conjugation is well-documented in numerous plasmid systems (76, 74, 9, 98, 75, 8). However, our results did not reveal any apparent fitness penalty associated with Sfx loss. *E. coli* carrying R6K or R6KΔ*sfx* display similar growth dynamics in nutrient-rich (LB) and poor (M9) environments (Figure 4B-E; Figure S4), and R6K*Δsfx* carriers exhibit growth defects only under the highly artificial condition in which chromosomal H-NS is also lost (Figure 4D). This lack of fitness cost is also found on a larger time scale, wherein R6KΔ*sfx* is stably maintained in the absence of antibiotic selection, consistent with previous studies on IncX3 plasmids (33), without selecting for mutations affecting conjugation efficiency (Figure 7). Our findings suggest that conjugatively derepressed plasmids may have a fitness advantage under certain conditions. Indeed, plasmids harboring “superspreader” mutations that increase conjugation efficiency have been isolated from clinical and environmental sources (99, 100). Our findings also imply that other selective forces, not measured in laboratory conditions, may impose a selective advantage to keeping plasmid conjugation rates low, such as susceptibility to bacteriophages that attach to conjugal pili (11–14).

In summary, our study provides insights into the molecular evolution and functionality of plasmid-encoded H-NS homologs. We demonstrate the evolutionary and functional divergence of Sfx from chromosomal H-NS and its unique role in plasmid conjugation repression. The altered protein-protein and protein-DNA interaction interfaces of Sfx, driven partially by positive selection, likely contribute to its distinct functionality. Despite the conservation of Sfx among IncX plasmids, its loss does not impose an obvious fitness cost, reflecting the complexity of selection forces shaping plasmid evolution. Our results, therefore, highlight the sequence and functional diversity within the H-NS family and underscore the pivotal role that protein evolution plays in plasmid biology.

## Methods and Materials

### Strains and plasmids

The strains, plasmids, and oligonucleotides used in this study are listed in Table S4, Table S5, and Table S6, respectively. *E. coli* DH5α (NEB) is used for cloning and plasmid propagation. Except for the bacterial-two-hybrid assays, all experiments are performed using the *E. coli* BW25113 or strains from the Keio collection when specified (101). For all conjugation assays, the recipient strain is a spontaneous NaN_3_^r^ EcoR25 strain derived in-house. *E. coli* BTH101 [F-, *cya-99, araD139, galE15, galK16, rpsL1* (Str^r^), *hsdR2, mcrA1, mcrB1*] and T18/T25-fragment containing plasmid backbones (pKT25, pKNT25, pUT18, pUT18C) are a kind gift from Véronique Taylor (University of Toronto) and are used for the bacterial-two-hybrid assays. The domain-swapping, complementation, and bacterial-two-hybrid plasmids are constructed via Gibson assembly using PCR-amplified vector backbone and fragments from R6K or *E. coli* BW25113 genomic DNA following the manufacturer’s instructions. Mutations and small fragment substitutions in plasmids are introduced via site-directed-mutagenesis using a homebrew KLD enzyme mix. Plasmids are transformed into *E. coli* using electroporation or heat shock. R6KΔ*sfx* is a kind gift from Irina Artsimovitch (Ohio State University). All plasmid constructs are verified via Sanger or Nanopore sequencing (Plasmidsaurus).

### Media and culture conditions

*E. coli* strains are routinely grown in LB liquid media (BD Difco™ LB Broth Miller) or LB agar (BD Difco™ LB Agar Miller) at 37°C. Liquid cultures are grown with shaking (200 rpm). When required for plasmid maintenance, the growth media is supplemented with 100 µg/mL streptomycin, 50 µg/mL kanamycin, 100 µg/mL ampicillin, 20 µg/mL chloramphenicol, and/or 0.01% NaN_3_. For growth curve, biofilm, and serial passaging, strains are grown in M9 (BD Difco™ Bacto M9 Minimal Salts) + 0.2% glucose when specified.

### Filter-paper conjugation assay

For all experiments, one biological replicate of the recipient strain and two biological replicates of the donor strains are grown overnight in LB (recipient: supplemented with 0.01% NaN_3_; donor: supplemented with 100 µg/mL of streptomycin and 20 µg/mL chloramphenicol if harboring pHSG576 constructs) at 37°C with shaking (200 rpm). The conjugation efficiency on filter-paper is assessed as follows: 2 OD_600_U of donor and recipient (spontaneous NaN_3_^r^ EcoR25) overnight cultures are pelleted (4500 x g for 5 minutes at room temperature), washed with sterile phosphate-buffered saline (PBS), and combined at equal volume. 20 µL of the culture mixture is spotted on MF-Millipore™ Membrane Filter (0.22 µm pore size) overlaid on LB agar and incubated at 30°C for 3 hours. The bacterial spots are washed off by vortexing the filter-paper discs in 100 µL of sterile PBS for 20 seconds. The donor and transconjugant populations are determined by spotting serially diluted bacterial mixtures on LB agar supplemented with 100 µg/mL ampicillin and LB agar supplemented with 100 µg/mL ampicillin and 0.01% NaN_3_, respectively. Each conjugation experiment is repeated at least twice.

### Biofilm conjugation assay

Donor and recipient strains are grown and processed via the same method outlined in “Filter-paper conjugation assay.” For each strain, washed overnights of three biological replicates are diluted to a final OD_600_ of 0.03 in 150 µL of M9+0.2% glucose media in a 96-well polystyrene plate (Sarstedt). The similarity in donor cell input (for conjugation assays) or R6K/R6KΔ*sfx* carrier (for biofilm competition assay) is validated by plating serially diluted inoculum on LB agar supplemented with 100 µg/mL streptomycin (which selects for both R6K and R6KΔ*sfx* carriers) and/or 50 µg/mL kanamycin (which selects for only R6KΔ*sfx* carriers.). Biofilm formation is allowed to occur statically at 30°C for 24 hours. After 24 hours, the planktonic cultures are aspirated and quantified via selective plating (see below for conditions). The biofilm is gently washed once with 150 µL of sterile PBS to remove non-adherent cells and resuspended in 40 µL of sterile PBS. To assess donor, recipient, and transconjugant population, the planktonic cultures/resuspended biofilms are serial diluted and spotted on LB agar supplemented with 100 µg/mL ampicillin, or 0.01% NaN_3_, or 100 µg/mL ampicillin and 0.01% NaN_3_, respectively. Each biofilm conjugation experiment is repeated at least twice.

### Bacterial-two-hybrid assay

The coding sequences of *hns, stpA*, *hha*, and *cnu* are amplified via PCR from *E. coli* BW25113 genomic DNA, whereas the coding sequences of *sfx* is amplified from R6K. The PCR products are cloned into bacterial-two-hybrid vectors (pUT18, pUT18C, pKT25, pKNT25) using Gibson assembly following the manufacturer’s instructions and validated by Sanger Sequencing. All H-NS homolog constructs (Sfx, H-NS, and StpA) are fused to the adenylate cyclase fragment at its C-terminus to preserve the N-terminal dimerization domain. High-copy vectors (pUT18C) encoding for Hha and Cnu are constructed with only the N-terminal T18 fusion (given that C-terminal fusion disrupts H-NS-Hha interaction (72)), whereas both N-terminal and C-terminal chimeric proteins are constructed for low-copy vectors. To detect protein-protein interactions, we heat-shocked the bacterial-two-hybrid vectors into *E. coli* BTH101 and plated them on LB supplemented with 100 µg/mL ampicillin and 50 µg/mL kanamycin to select for transformants. Three biological replicates of each strain are grown overnight in LB supplemented with 100 µg/mL ampicillin and 50 µg/mL kanamycin at 37°C with shaking (200 rpm). Expression of the chimeric proteins is then induced by spotting 5 µL of overnight cultures on LB agar supplemented with 100 µg/mL ampicillin, 50 µg/mL kanamycin, 40 µg/mL X-gal, and 50 mM IPTG. The plates are incubated at 30°C for 24 hours (for strains carrying pUT18::*hns* (M1-G85), pUT18::*stpA,* pUT18::*sfx,* pUT18::*sfx-*N-truncated) or 48 hours (for strains carrying pUT18C::*hha* and pUT18C::*cnu*). Each bacterial-two-hybrid screen is repeated twice.

### Serial passaging and assessing passaged strain conjugation efficiency

6 independent lineages of *E. coli* carrying R6K or R6KΔ*sfx* are serially passaged in LB or M9+0.2% glucose for 20 days (∼200 generations). All strains are grown in 5 mL of media in 20 mm glass tubes at 37°C with shaking (200 rpm). Every 24 hours, the cultures are diluted 1:1000X in fresh media. The proportion of plasmid-retaining cells is assessed by plating serially diluted cultures on LB agar (quantify total bacterial population) and LB agar supplemented with 100 µg/mL streptomycin (which selects for only plasmid carriers). To assess whether mutations accumulated on the R6K/R6KΔ*sfx* plasmid during passaging, single colonies of passaged strain (from frozen DMSO stocks) were grown overnight in LB supplemented with 100 µg/mL streptomycin. Plasmids were harvested using the GeneJET Plasmid Miniprep Kit following the manufacturer’s protocol and sequenced using Nanopore sequencing (Plasmidsaurus). The effect of serial passaging on plasmid conjugation is assessed by taking single colonies from day 0 (ancestral strain) and day 20 (passaged strain) DMSO stocks preserved at -80°C and evaluating the filter-paper conjugation efficiency of two biological replicates, each with two technical replicates (filter-paper conjugation experiment is performed as specified above).

### Growth curve analysis

1 OD_600_U of overnight *E. coli* cultures are pelleted (4500 x g for 5 minutes at room temperature), washed with sterile PBS, and diluted to a final OD_600_ of 0.02 in 200 µL of growth media (LB or M9+0.2% glucose supplemented with 20 µg/mL chloramphenicol if strains carry pHSG576 constructs). All cells are seeded in 96-well polystyrene plates (Sarstedt), sealed with Breathe-Easy® sealing membrane (Sigma Aldrich), and grown at 37°C for 24 hours with shaking every 15 minutes. OD_620_ is monitored every 15 minutes by an S&P growth robot. All growth curve assays are repeated at least three times.

### Motility assay

Overnight *E. coli* cultures grown at 37°C with shaking are subcultured 1:100X in 5 mL of LB with no antibiotics to an OD_600_ of ∼0.6. 6 µL of mid-log culture is aspirated into the middle of 0.3% LB agar, dried at room temperature for 1 hour, and incubated at 30°C for 24 hours. The motility assay is repeated twice.

### Conservation and alignment of chromosomal and plasmid-borne H-NS homologs

We aligned the amino acid sequences of Sfx (NCBI accession ID: WP_001282381.1), Acr2 (NCBI accession ID: WP_000651490.1), StpA (Uniprot accession: P0ACG1), Sfh (Uniprot accession: Q8GKU0), and H-NS (Uniprot accession: P0ACF8) using MAFFT G-INS-i (102). The conservation within Sfx homologs is derived using the ConSurf webserver (103) with default parameters.

### Generating Sfx, H-NS, and StpA sequence alignment for molecular evolution analysis

The NCBI nr/nt database (104) is queried using tblastn in October 2022 with default search parameters to retrieve nucleotide sequence homologous to Sfx from the R6K plasmid (NCBI accession ID: WP_001282381.1), H-NS from *E. coli* K12 (Uniprot ID: P0ACF8), and StpA from *E. coli* K12 (Uniprot ID: P0ACG1). We removed all hits that are <50% similar to the seed template sequences and retrieved the coding sequences corresponding to the remaining hits using an in-house script. The resulting sequences are filtered to eliminate short (<88 amino acids for Sfx homologs, <100 amino acids for H-NS and StpA homologs) and overly long peptides (>200 amino acids for all sequences) that are indicative of pseudogenes and/or misannotation. The nucleotide sequences are clustered using MMseqs2 (105) to identify sequence-level representatives (95% identity, 80% coverage, coverage mode=1). The sequence representatives are aligned using the Guidance2 web server (106) (codon model, MAFFT with maximum 1000 cycles of iterations) and trimmed to eliminate any sequences with <0.6 confidence score and columns with <0.87 confidence score. To aid with phylogeny visualization, the Ler protein from *E. coli* O157:H7 str. Sakai (Genbank accession: BAB38011.2), a H-NS-like protein that is distantly related to Sfx, H-NS, and StpA ((107); pairwise sequence identity ∼20-25%), is chosen as the outgroup and aligned with all sequences.

### Constructing phylogeny of Sfx, H-NS, and StpA

A Sfx, H-NS, and StpA phylogeny is inferred via the maximum-likelihood method using IQ-TREE2 (108). The best codon substitution model is chosen using ModelFinder (109), and branch confidence is assessed via 5000 rounds of Ultrafast Bootstrap Approximation (110) and 1000 bootstrap replicates of SH-aLRT (111). For visualization purposes, the tree is rooted at the Ler protein, but all molecular evolution analyses are performed on an unrooted phylogeny that does not contain the Ler protein sequence. Sequence taxonomy is derived based on the subfamily classification proposed by Alnajar and Gupta (112), while other sequence characteristics (genetic context, AT-hook motif) are retrieved from NCBI or derived from the protein alignment and mapped onto the phylogeny using the *ggtree* package (113).

### Molecular evolution analysis using PAML

We used PAML 4.10.6 (114) to explore the variation in selective constraints across Sfx, H-NS, and StpA homologs. Branch-sites (115) and Clade models (116) are fitted using the codeml program, and likelihood ratio tests are used to explore whether using more complex models leads to a statistically significant improvement in model fit (117). We used the Akaike information criterion (AIC) as an evaluation metric for comparison across non-nested models, as suggested by Weadick and Chang (117). Residues under positive selection are detected using NEB and BEB analyses (44).

### Constructing WebLogo of Sfx, H-NS, and StpA homologs

Sequences within the Sfx, H-NS, and StpA clades (defined based on phylogeny) are aligned (within-clade) using the MAFFT E-INS-i algorithm (102). The sequence alignments are manually trimmed using Pfaat to remove gappy regions (118), and the amino acid frequency at each position is visualized using the WebLogo webserver (119).

### Mapping Sfx orthologs across IncX plasmids

975 IncX plasmids, as classified by PlasmidFinder (120), were retrieved from PLSDB (121) in March of 2023. One plasmid is arbitrarily selected from each unique PlasmidFinder lineage to produce 55 representative sequences. For each lineage representative, we reannotated the sequences using PGAP (122) and derived plasmid attributes (Replicon, relaxase, MPF, OriT, predicted mobility, predicted host range) using MOB-suite (123). To identify H-NS orthologs and co-occurring proteins, we performed orthologous clustering of all 55 lineages using OrthoFinder with default parameters (124). The AT-hook motif of the H-NS homologs are derived by performing multiple sequence alignment using the DECIPHER package (125) and further validated by visual inspection. The attributes of the representative plasmids are clustered via hierarchical clustering using Gower distance and visualized using the R package *ggtree* (113).

### AlphaFold structural prediction of Sfx, H-NS, and StpA

Tetramers and dimers of Sfx, H-NS, and StpA are predicted using ColabFold (Alphafold2-multimer) (126). Template information of each sequence set is fetched from the PDB70 database, and the predicted structures are relaxed using the amber force fields. The secondary structure is derived using POLYVIEW-2D (127) and visualized using the R package *gggenes* (128), while the 3D structure is modeled in PyMOL (129).

### GC content calculation of IS and non-IS regions in IncX plasmids

22 IncX plasmid representatives that carry H-NS homologs with the RGR AT-hook motif are searched for putative IS regions using ISfinder (blastn, default parameters) (130). The resulting hits are filtered to retain only higher confidence (e-value ≤ 0.1) and larger sequences (>100 bp). An in-house script is then used to calculate the GC% of all putative IS and non-IS regions.

## Acknowledgments

This work was funded by grants from the Natural Sciences and Engineering Research Council of Canada (NSERC: RGPIN-2020-06015 and USRA 24/25). We thank members of the Navarre Lab, Jordan Lin (University of Toronto), Dr. Irina Artsimovitch (Ohio State University), Dr. Barbara Funnell (University of Toronto), and Dr. Kamna Singh for their feedback on the manuscript and technical support. We would also like to thank Dr. Véronique Taylor (University of Toronto) for her help in setting up the bacterial-two-hybrid assay and sharing the strains and plasmids. Lastly, we thank Dr. Belinda Chang for her guidance and critical feedback on the molecular evolution analysis.

## Supplementary Figure Legends

**Figure S1. MPF_T_-type IncX plasmids frequently carry Sfx homologs.** 55 representative IncX lineages were analyzed using orthologous clustering and MOB-Typer to identify H-NS homologs and characterize plasmid attributes. The plasmids are clustered using hierarchical clustering based on various attributes, such as the AT-hook motif of the H-NS homolog (“H-NS"), predicted replicon type (“Replicon”), MPF (mating pair formation) type, predicted mobility (“Mobility”), predicted host range (“Range”), and GC%. Note that plasmids with a H-NS label of “RGR and QGR” carry two H-NS homologs, one with a RGR AT-hook motif and the other with a QGR AT-hook motif. The node representing R6K is marked with a blue diamond.

**Figure S2. The StpA clade exhibits less selective constraint.** The gene map is colored based on the site classification derived from Bayes Empirical Bayes (BEB) analysis of the best performing multi-clade CmD alternative model (Sfx and StpA clades set as foregrounds). 3 site classes are listed, each representing a specific ratio of nonsynonymous substitution rates to synonymous substitution rates (ω). Blue squares represent sites under the highest degree of purifying selection followed by the magenta squares (neutrally evolving sites). The yellow squares represent sites under different selective constraints across the Sfx, StpA, and H-NS clades. The corresponding secondary structure at each alignment position is shown above the heatmap and is derived from a Colabfold-predicted structure of *E. coli* K12 H-NS (this is used instead of Sfx due to gaps in the alignment of the linker region). Abbreviations: α (alpha-helix); β (beta-sheet).

**Figure S3. Positively selected residue positions are mostly conserved within phylogenetic clades.** Sequences within the Sfx, H-NS, and StpA clades (See Figure 1 for details) are aligned using MAFFT E-INS-i. The conservation of amino acids at each position is visualized using WebLogo. The locations of positively selected sites, as calculated by Naive Empirical Bayes and Bayes Empirical Bayes analysis, are highlighted in yellow. The amino acid transitions from Sfx (R6K) to *E. coli* K12 H-NS are printed above each position.

**Figure S4. Sfx loss does not affect R6K carrier growth in biofilms.** (A) Sfx loss does not affect R6K donor growth in a biofilm conjugation setting. Plotted on the y-axis are the final donor population densities of R6K (left) and R6KΔ*sfx* donors in the biofilm (left panel) and planktonic (right panel) states. Briefly, R6K/R6KΔ*sfx* donors and EcoR25 recipients (NaN_3_^R^) are seeded at a starting OD_600_ of 0.03 and cultured statically at 30°C in 150 µL of M9+0.2% glucose media in a 96-well polystyrene plate. The final population density of donor, transconjugant, and recipient cells after 24 hours is assessed by selective plating. Statistical significance is assessed using the Student’s t-test with Benjamini-Hochberg correction. (B) R6K and R6KΔ*sfx* carriers have similar growth rates in a biofilm setting. Each jointed point represents the final population density of R6K (left) and R6KΔ*sfx* (right) carriers in the biofilm (left panel) and planktonic (right panel) states. The competition assay is performed by inoculating R6K and R6KΔ*sfx* carriers to a final OD_600_ of 0.03 and coculturing them at 30°C in 150 µL of M9+0.2% glucose media in a 96-well polystyrene plate. The final population composition of R6K and R6KΔ*sfx* carriers after 24 hours of incubation is assessed using selective plating. Statistical significance is assessed using the Wilcoxon matched pairs test. (A-B) displays the pooled results from two experimental replicates, with each point representing a biological replicate (averaged value across two technical replicates). Abbreviations: ns (adjusted *p-*value ≥ 0.05).

**Figure S5. IncX plasmids carrying Sfx homologs display an atypical base composition.** (A) R6K cargo genes have a higher GC% than core plasmid genes. The line plot displays the 2kbp sliding-window average GC% across the R6K sequence. The corresponding gene annotation derived from the RefSeq database (NCBI accession: NZ_LT827129.1) is shown below. Genes are colored based on functional categorization. The average GC% of H-NS binding sites (38.6% GC) is indicated by a blue dashed line. (B) IncX plasmids with Sfx homologs display an atypical base composition. The paired dot plot displays the GC% of non-insertion element (non-IS) and insertion element (IS) regions of 19 IncX plasmid representatives that carry a Sfx homolog. Each connected pair of points represents the value of one plasmid. The points representing R6K are colored in navy blue.

